# Programs, Origins, and Niches of Immunomodulatory Myeloid Cells in Gliomas

**DOI:** 10.1101/2023.10.24.563466

**Authors:** Tyler E. Miller, Chadi A. El Farran, Charles P. Couturier, Zeyu Chen, Joshua P. D’Antonio, Julia Verga, Martin A. Villanueva, L. Nicolas Gonzalez Castro, Yuzhou Evelyn Tong, Tariq Al Saadi, Andrew N. Chiocca, David S. Fischer, Dieter Henrik Heiland, Jennifer L. Guerriero, Kevin Petrecca, Mario L. Suva, Alex K. Shalek, Bradley E. Bernstein

## Abstract

Gliomas are incurable malignancies notable for an immunosuppressive microenvironment with abundant myeloid cells whose immunomodulatory properties remain poorly defined. Here, utilizing scRNA-seq data for 183,062 myeloid cells from 85 human tumors, we discover that nearly all glioma-associated myeloid cells express at least one of four immunomodulatory activity programs: Scavenger Immunosuppressive, C1Q Immunosuppressive, CXCR4 Inflammatory, and IL1B Inflammatory. All four programs are present in IDH1 mutant and wild-type gliomas and are expressed in macrophages, monocytes, and microglia whether of blood or resident myeloid cell origins. Integrating our scRNA-seq data with mitochondrial DNA-based lineage tracing, spatial transcriptomics, and organoid explant systems that model peripheral monocyte infiltration, we show that these programs are driven by microenvironmental cues and therapies rather than myeloid cell type, origin, or mutation status. The C1Q Immunosuppressive program is driven by routinely administered dexamethasone. The Scavenger Immunosuppressive program includes ligands with established roles in T-cell suppression, is induced in hypoxic regions, and is associated with immunotherapy resistance. Both immunosuppressive programs are less prevalent in lower-grade gliomas, which are instead enriched for the CXCR4 Inflammatory program. Our study provides a framework to understand immunomodulatory myeloid cells in glioma, and a foundation to develop more effective immunotherapies.

## INTRODUCTION

Diffuse gliomas are the most common primary malignant brain tumors in adults, and remain ultimately fatal despite significant advances in our molecular understanding of the malignant cells^1–7^. These tumors are divided into isocitrate dehydrogenase (IDH)-mutant and wild-type (WT) gliomas^8^, with glioblastoma (GBM), IDH-WT, being the most prevalent and aggressive form (median overall survival <2 years)^9,10^. The limited efficacy of current therapies, which include surgery, chemotherapy, and radiotherapy^11^, underscores the need for novel therapeutic strategies.

Immunotherapy has revolutionized treatment for many types of cancer. Unfortunately, despite anecdotal responses^12,13^, immunotherapy trials have failed to provide life-prolonging benefit for glioma patients^14,15^. Gliomas represent an immunotherapy challenge due to the unique immune microenvironment of the brain, restricted access of systemic therapies due to the blood-brain barrier, and the need to balance therapeutic immune responses with potentially fatal inflammation-induced edema. The poor clinical responses to conventional immunotherapy highlight the need to better understand the complex microenvironment in gliomas, which includes limited activated T-cells and an abundance of myeloid cells.

Tumor-associated myeloid cells have become a major focus in the pursuit of effective immunotherapies for solid tumors. In many solid tumors, including glioma, increased myeloid cells are associated with higher grade and worse overall survival^16,17^. These cells can create an immunosuppressive microenvironment that leads to immunotherapy resistance. Understanding their functional phenotypes, origins, and developmental drivers is a critical step towards rational therapeutic strategies that overcome myeloid immunosuppression.

In gliomas, myeloid cells are particularly suppressive and are the most prevalent non-malignant cell type, comprising up to 50% of all cells in a tumor^18^. Their abundance and ability to orchestrate neighboring cell behavior makes them central to the pathobiology of gliomas^19^. Prior studies have shown that myeloid cells have a major influence on the molecular state of tumor cells^6,20^, as well as tumor-infiltrating T cells^20–22^, the main effector cells of checkpoint blockade, vaccine, and chimeric antigen receptor (CAR)-T cell therapies. Tumor-associated myeloid cells also recruit additional myeloid cells from the peripheral circulation through cytokine and chemokine release, and may drive them towards immunosuppressive phenotypes^19^. However, the specific myeloid cell types and gene expression programs that orchestrate these functions remain to be determined.

Myeloid cells in gliomas have traditionally been classified and studied according to cell type and/or presumed developmental origin^4,18,21,23–25^. Myeloid cell types include microglia, macrophages, monocytes, conventional dendritic cells (cDC), and neutrophils. Origin is typically classified as microglia-derived or bone marrow-derived based on marker genes identified from lineage tracing experiments in healthy mice. These murine lineage tracing studies have shown that microglia are derived from the embryonic yolk sac and remain isolated to the brain, while other myeloid cell types are derived from bone marrow^26–28^. However, despite its therapeutic implications, the origins of myeloid cells in human brain tumors remain uncertain^29,30^.

Our understanding of the heterogeneity of malignant cells in glioma has greatly improved due to single-cell RNA sequencing (scRNA-seq) technologies. Over the past decade, this work has helped reveal the developmental origins and inherent plasticity of these cells, yielding insights into the function of the main cellular states (NPC-like, OPC-like, AC-like, MES1, and MES2) and suggesting rational targets to limit their progression^1,3–5,7^. Recent studies using various single-cell technologies have begun to uncover the diversity of myeloid cell states in human and mouse gliomas, including some interactions with other cell types within the tumors^4,18,20,21,23–25,31,32^. These studies revealed differences in the composition and suspected origin of myeloid cell types between IDH-mutant and wild type gliomas, primary and recurrent gliomas, and even within different regions of the same tumor^21,24^.

Yet, many outstanding questions remain. First, at present, there is no consensus on the definition of myeloid cell states, or how they inform the clinical and biological features of gliomas. Second, previous studies have viewed myeloid cells through the lens of the traditional cell type and origin classification, but classifying functional activities independent of cell type or origin has been challenging with standard cell clustering approaches. Third, the origins of myeloid cells in gliomas remain uncertain due to difficulties in tracing cell lineage in human samples. Finally, the interplay between myeloid cells and other malignant and non-malignant cell states within the tumor has primarily been deduced from variations in cellular composition within samples or with limited markers. Assessing the spatial relationships of these cells at increased granularity is crucial to understand how myeloid cells interact with their niches and immune microenvironments. Thus, incomplete knowledge of glioma-associated myeloid cells, their diverse expression programs, their origins, and their functional significance within the specialized glioma immune microenvironment remains a major impediment to advancing immune therapies.

An additional impediment relates to challenges with experimental modeling of human tumor-associated myeloid cells. Tumor-associated macrophages change state quickly *in vitro* on monolayer plastic cell culture, and mouse models incompletely recapitulate macrophage programs associated with human tumors^33^. Mouse microglia are smaller, are less morphologically complex than their human counterparts, and lack orthologues to important human microglial genes^34^. While these systems have helped answer important questions and demonstrated the importance of myeloid cells for glioma biology^20,31,35^, more faithful and representative experimental systems of human tumor-associated myeloid cells are urgently needed for both fundamental understanding and clinical translation.

Here we sought to overcome these limitations through a systematic single-cell study of myeloid cells in human gliomas coupled with functional validations in refined experimental tumor models. We leveraged scRNA-seq data for 85 diverse gliomas, including primary and recurrent IDH-mutant and wild-type tumors, and emerging computational methods for decoupling myeloid cell type from activity to identify four dominant immunomodulatory activity programs shared across microglia, macrophages, monocytes, and dendritic cells. We then integrated lineage tracing techniques in patient samples, spatial transcriptomics, and high-fidelity *ex vivo* human tumor models to discover the cellular origins, tumor niches, and drivers of these dominant immunomodulatory programs. Our analyses portray a dynamic and plastic myeloid cell compartment that is responsive to microenvironmental cues and evolves with glioma progression to become highly immunosuppressive. In sum, they provide a foundation for advancing diagnostic and immunotherapeutic strategies for gliomas.

## RESULTS

### Unbiased identification of consensus gene programs in glioma-associated myeloid cells

To better understand the immune microenvironment in gliomas, we utilized scRNA-seq to characterize all immune and non-immune cell types within freshly resected human adult diffuse gliomas. We included a wide array of tumors, spanning IDH-wild type and mutant tumors, primary and recurrent tumors, and tumors exposed to different therapies. We combined 43 tumor profiles prospectively collected for this study with an additional 42 consolidated from prior publications^7,21,36^. These 85 profiles (Supplemental Table 1) were divided into a discovery dataset that included 44 tumors profiled by the latest 3’ scRNA-seq technologies, (10Xv3 / SeqWell S^3^)^37^, and a validation dataset (41 tumors profiled by 10Xv2). We annotated all cells based on marker gene expression, removed doublets, and called single-cell copy number alterations (CNAs) to confirm malignant cells (Fig. 1a, Extended Data Fig. 1a-c, see Methods).

**Fig. 1:**
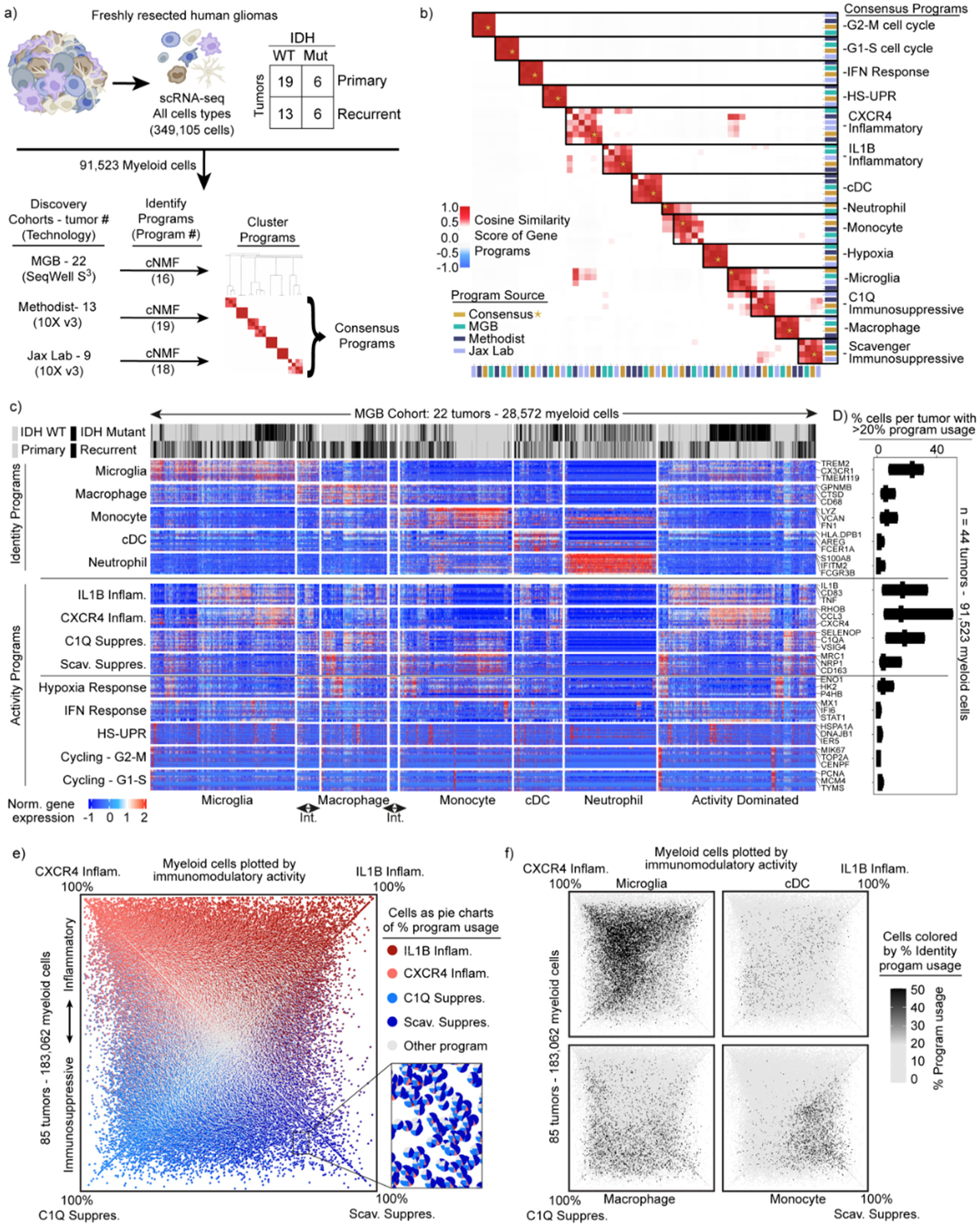
Identification of consensus superimposable myeloid cell identity and cell activity programs. a) Schematics of the analysis pipeline for identifying the recurrent myeloid programs across the three discovery glioma cohorts. b) Heatmap demonstrating the cosine similarity indices of the gene spectra scores of each program in the three discovery cohorts. c) Heatmaps demonstrating the expression of genes in recurrent myeloid programs (rows) by cell (column) grouped by cell type. Cell type defined by usage of myeloid identity programs. d) Box plots exhibiting the percentage of cells by sample expressing the recurrent myeloid programs indicated on the left of the heatmap across the three discovery cohorts. Int. = intermediate cells expressing adjacent identity programs. e) Quadrant plot of myeloid cells from the discovery and validation cohorts. Coordinates of cell determined by: (CXCR4 - Scav. program usage), (IL1B - C1Q program usage). Axes are diagonal. Each dot is a pie chart exhibiting the prevalence of the four indicated immunomodulatory programs in that cell. f) Quadrant plots displaying myeloid cell identify usage per cell.

We then turned our attention to the myeloid cells. To discover the consensus myeloid gene expression programs in gliomas, we utilized our discovery dataset, which was composed of three independent cohorts from three different institutions. We used an unbiased method, consensus non-negative matrix factorization (cNMF)^38^ to identify sets of genes (‘programs’) that were coordinately regulated across the myeloid cells within each cohort (Supplemental Table 2, Methods). Hierarchical clustering of these programs identified recurrent expression programs captured in all three discovery cohorts, from which we derived 14 consensus gene programs (Fig. 1a, Extended Data 1a-c). These 14 programs captured the gene expression patterns of all individual programs within the corresponding clusters (Fig. 1b).

These consensus programs included both myeloid cell identity programs and cell activity programs. The identity programs contain classical marker genes for myeloid cell types, including microglia, macrophages, monocytes, dendritic cells, and neutrophils. The activity programs are composed of genes with immunomodulatory functions, genes involved in specific cell response programs (such as interferon or hypoxia response) and genes linked to proliferation (Fig. 1c). The 14 programs were found across IDH-mutant, IDH WT, primary, and recurrent gliomas (Fig. 1c), and importantly, were all recapitulated in our validation cohort (Extended Data Fig. 1d-e).

In parallel, we performed Louvain clustering and Uniform Manifold Approximation and Projection (UMAP) to understand the myeloid cell state spaces. This standard approach treats cells as a singular unit, clustering them based on their similarity to other cells, as opposed to cNMF which considers multiple discrete programs in each single cell by computing and evaluating the usage of consensus gene programs. The clustering and UMAP visualization highlighted different myeloid cell types seen in our cNMF analysis, but was less effective at capturing the cNMF activity programs, each of which can manifest in different cell types (Extended Data Fig. 2a-c). Going forward, we relied on cNMF to evaluate myeloid cell types and their superimposed activity programs, given its ability to capture more than one program in a given cell.

### Superimposable myeloid cell identity and cell activity programs

Among the five cell identity programs, we find that the microglia program, highlighted by classical marker genes *TMEM119*, *P2RY12*, and *CX3CR1*, is the most prevalent (Fig. 1d). The macrophage program includes *GPNMB*, *LGALS3*, *CD63*, *CD9*, and *CD68*, all well-established markers of tumor-associated macrophages. The cDC program, which is composed of cDC1 and cDC2 marker genes, was the least prevalent. While the neutrophil and cDC programs were almost entirely composed of known peripheral neutrophil and cDC genes, the tumor-associated monocyte program had significant differences from peripheral monocytes. To investigate this further, we performed scRNA-seq on peripheral myeloid cells from matched blood samples for 17 of our patients (Extended Data Fig. 3). cNMF analysis of the peripheral cells identified three monocyte programs (CD14, CD16 and Suppressive; see Methods). We found that the tumor-associated monocyte program shared features with the CD14 and Suppressive programs in peripheral monocytes, but had almost no overlap with the CD16 monocyte program (Extended Data Fig. 3). The tumor-associated monocyte program also included genes involved in cell adhesion, migration, differentiation, and initial inflammatory response (e.g., *VCAN, FCN1, LYZ, CD44, FLNA*, and *CCR2*). This suggests that the monocytes represented in the GBM data are undergoing differentiation within the tumor tissue.

Notably, the most prevalent programs in the glioma-associated myeloid cells were four activity programs enriched for immunomodulatory genes (Fig. 1d and Extended Data Fig. 1e). 91% of the myeloid cells expressed one of these four immunomodulatory programs. For comparison, roughly 76% of cells could be confidently assigned to one of the five myeloid cell types based on the cNMF identity programs (the other 24% were ‘activity dominated’) (Fig. 1c). The immunomodulatory programs could be split into two inflammatory programs and two immunosuppressive programs based on the genes driving the programs (Fig. 1b-c). The IL1B Inflammatory program includes inflammatory cytokines and chemokines with established roles in myeloid cell recruitment such as *IL1B, IL1A, CCL3, CCL4, CC2, TNF, OSM*, and *CXCL8*. The CXCR4 Inflammatory program is composed of genes involved in lymphocyte and monocyte recruitment such as *CXCR4, CXCL12, CCL3, CCL4*, and *CX3CR1*, as well as genes involved in immediate stress responses like *RHOB, JUN, KLF2*, and *EGR1*. Interestingly, this program also includes genes known to interact with neural cell types, such as *PDK4, P2RY13*, and *CXCR4*. On the immunosuppressive side, the C1Q program is defined by expression of *C1QA, C1QB, C1QC, CD16, CD163, C3, C2,* and *VSIG4*, many of which are involved in the complement system and/or have established immunosuppressive effects in other contexts. Finally, the Scavenger Immunosuppressive program is composed of scavenger receptors, such as *MRC1, MSR1, CD163, LYVE1, COLEC12* and *STAB1*, along with other potentially immunosuppressive genes such as *NRP1, RNASE1* and *CTSB*. Many of these have been shown to suppress T cell function including CD163^39^ and VISG4^40^ which bind to T cells and inhibits their proliferation, as well as MSR1 (CD204) which has a soluble form that binds and inhibits IFN-γ from activating T cells through inhibiting STAT1 signaling^41^.

Each of the four programs are expressed in multiple cell types; for example, the IL1B Inflammatory program is found in subsets of all myeloid cell types (Extended Data Fig. 4a-b). Conversely, each myeloid cell type utilizes more than one of the four immunomodulatory activity programs. Macrophages can express any of the four programs, but are enriched in the two immunosuppressive programs (Extended Data Fig. 4c). Microglia are enriched for the inflammatory programs and the C1Q Immunosuppressive program, but rarely express the Scavenger Immunosuppressive program. Neutrophils are unique in that they have limited expression of the four immunomodulatory programs, but rather are typically dominated by the neutrophil program itself.

These findings prompted us to seek more holistic insight into the four immunomodulatory programs, their inter-relationships, and their usages across cell types. We plotted all 183,062 myeloid cells from the 85 tumors by their usage of each program (Extended Data Fig. 4d-e, Fig. 1e). Although the activity programs are driven by different sets of genes, they can be co-expressed within individual myeloid cells. The integrative analysis also revealed correlations (and anti-correlations) in the expression or ‘usage’ of the different activity programs across cells, while also affirming their associations with the different myeloid cell types (Fig. 1f, Extended Data Fig. 4c). Importantly, these patterns and distributions were conserved across all three discovery cohorts and the validation cohort (Extended Data Fig. 4f).

These collective analyses revealed four immunomodulatory activity programs utilized by the multiple myeloid cell types in human gliomas, and present regardless of IDH mutation, recurrence, or treatment status. Interestingly, only one of the four programs, the IL1B inflammatory program, was evident in peripheral myeloid cells in glioma patients (Jaccard Index > 0.1, Extended Data Fig. 3, Supplemental Table 2). This suggests that the myeloid cells are highly plastic and that their programs are directed by cell-extrinsic factors in the tumor microenvironment more than their origin.

### Convergent phenotypes of microglia- and bone marrow-derived myeloid cells in glioma

To gain further insight into the determinants and plasticity of myeloid cell phenotypes we investigated their cellular origins. The current paradigm based on mouse models is that microglia are self-renewing tissue resident macrophages derived from embryonic yolk sack, whereas other myeloid cell types, including immunosuppressive macrophages, come from bone marrow^24,26–28^.

Mitochondrial DNA mutations can be used as endogenous barcodes in human samples to infer lineage relationships and cellular origins. We utilized MAESTER^42^ to call mitochondrial DNA mutations in tumor-associated myeloid cells and matched peripheral blood monocytes from four patients (Fig. 2a). We distinguished mitochondrial mutations present in peripheral blood cells from those that were detected only in tumor-associated myeloid cells. We presumed that myeloid cells in the tumor whose variants matched the former were blood-derived, while those with the latter variants would likely represent resident myeloid cells in the brain.

**Fig. 2:**
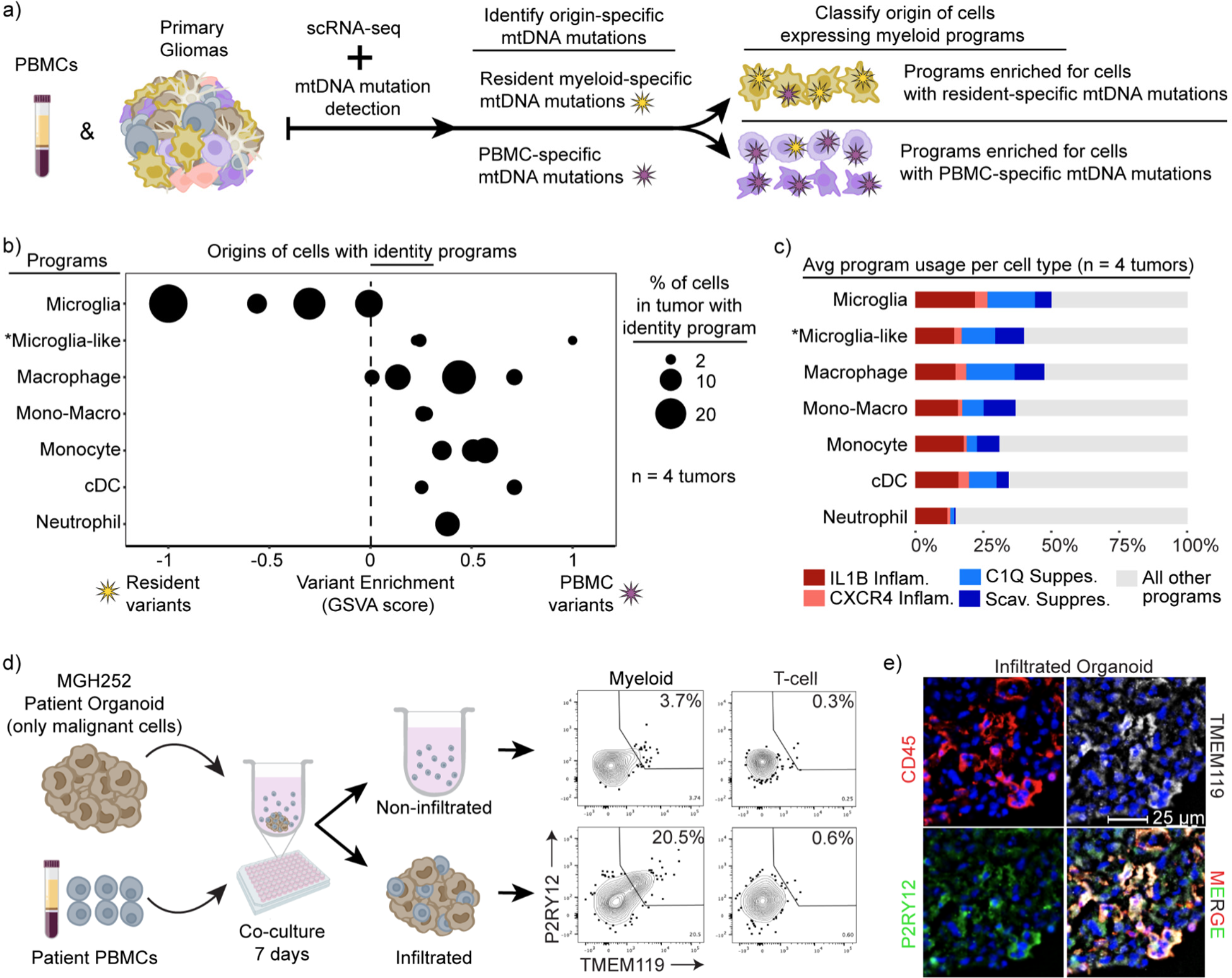
Convergent phenotypes of microglia- and bone marrow-derived myeloid cells in glioma. a) Schematics of the MAESTER analysis pipeline for determining the origin of myeloid cells in the glioma microenvironment. b) Dot plot exhibiting the enrichment difference between PBMC-specific and Resident-specific variants. Each dot represents the enrichment level of the indicated identities (left) in each patient. X-axis denotes the scaled difference between GSVA enrichment of PBMC-specific variants and Resident variants. c) Stacked bar charts indicating the average usage of the indicated myeloid programs in the key across the four patients. The “other programs” category encompasses the other identities and activities. d) Schematic (left) and flow cytometry plots (right) of myeloid cells from indicated condition. T cells are used as gating control for P2RY12 and TMEM119. e) Immunofluorescence image showing matched patient derived PBMC cells infiltrated into a glioblastoma organoid.

Consistent with expectations, we found that cells expressing a microglia program were most likely to harbor resident myeloid cell-specific variants, while other myeloid cell types were more likely to harbor variants shared with peripheral blood (Fig. 2b). The cell activity programs were more promiscuous in terms of origins, manifesting across different cell types and derivations (Fig. 2c). Notably, intermediate cells that co-express microglia and macrophage programs were also enriched for peripheral blood variants (Fig. 2b). This suggests that bone marrow-derived myeloid cells can activate a microglia-like phenotype in tumors.

This result prompted us to directly evaluate the capacity of bone marrow-derived cells to acquire these glioma-associated myeloid phenotypes. We applied patient peripheral blood mononuclear cells (PBMCs) to glioma organoids derived from the same patient’s tumor resection that no longer contained immune cells (Fig. 2d). After one week of co-culture, we found that the organoids were extensively infiltrated by myeloid cells. We extracted these infiltrated myeloid cells and compared them to myeloid cells that remained in the surrounding media by flow cytometry. We found that the infiltrating myeloid cells up-regulated the canonical microglia markers, TMEM119 and P2RY12, confirming that bone-marrow derived monocytes can acquire features of the tissue resident microglia (Fig. 2d). In contrast, myeloid cells that remained outside the organoids were much less likely to express these markers. Immunohistochemistry of the organoids confirmed robust infiltration of immune cells, including myeloid cells expressing both microglia markers (Fig. 2e and Extended Data 5a-b). Interestingly, IFN-γ markedly increased infiltration and differentiation of myeloid cells applied to the organoids, consistent with prior work^43^ (Extended Data Fig. 5b-c).

Together these data show that all tumor-associated myeloid programs, including the microglia program, can be expressed in cells derived from the peripheral blood. They highlight the plasticity of myeloid cells in the tumor microenvironment and demonstrate that developmental origin does not constrain the expression of immunomodulatory programs in glioma-associated myeloid cells.

### Immunomodulatory program composition varies with histopathological tumor grade

We next asked whether the myeloid cell identities and immunomodulatory programs correlate with clinical factors such as IDH mutation status. Prior studies have reported increased inflammatory phenotypes in IDH-mutant tumors^18,25,32^. Consistently, we found that IDH-mutant tumors have a distinct composition of immunomodulatory myeloid programs, characterized by strong enrichment of the CXCR4 Inflammatory program and depletion of both immunosuppressive programs (Fig. 3a and Extended Data Fig. 6a-b). Prior studies have also reported increased microglia in IDH-mutant tumors^4,18,25,32^. However, we did not detect any significant difference in the composition of cell identity programs in our datasets, with the exception that the monocyte program was more common in IDH-WT tumors. Although the CXCR4 program manifests across multiple myeloid cell types, it has some overlapping markers with microglia that could make it difficult to distinguish from microglia using technologies that rely on limited marker genes.

**Fig. 3:**
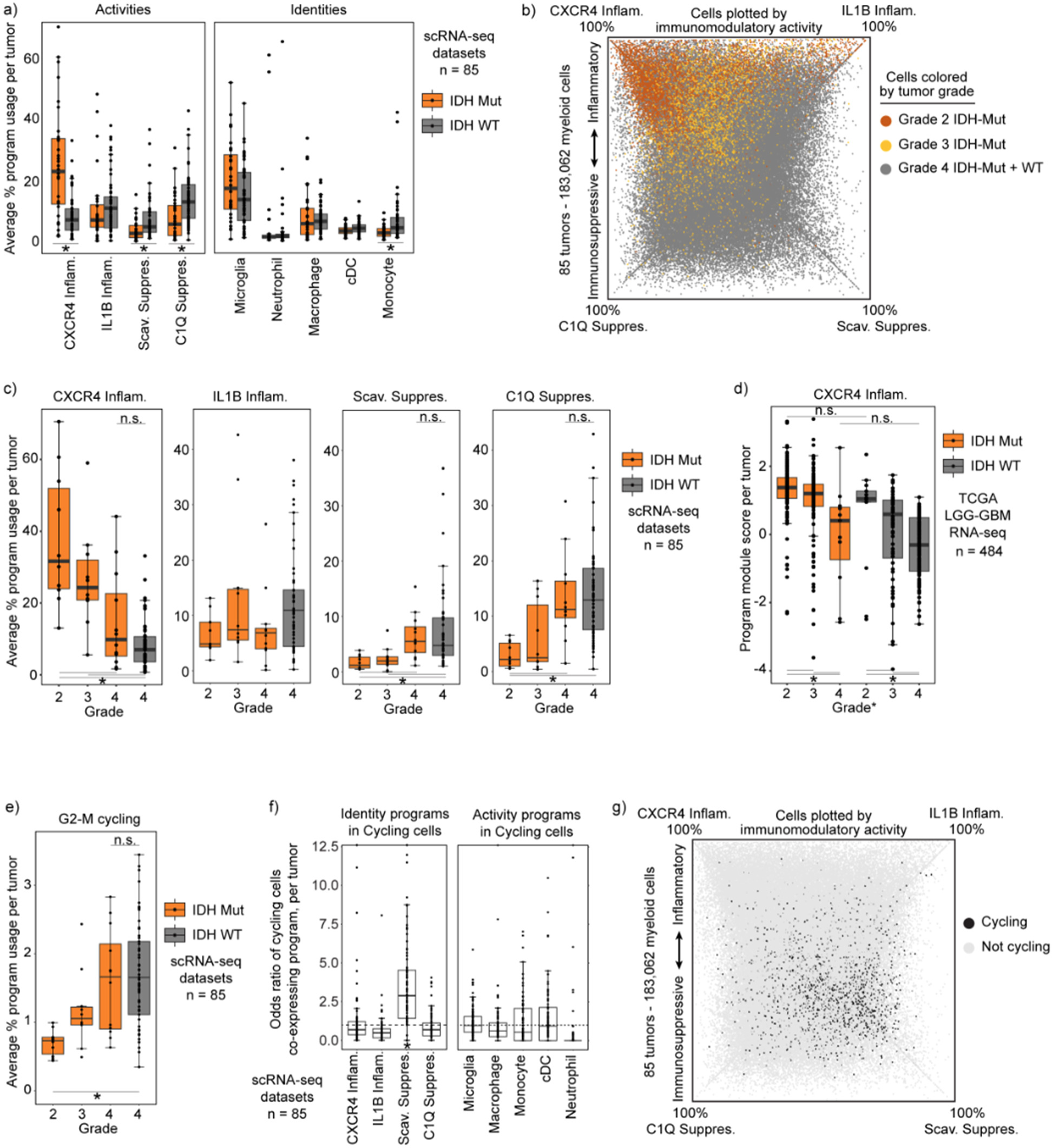
Immunomodulatory program composition varies with histopathological tumor grade. a) Boxplot exhibiting the average usage of the indicated activity or identity programs. *FDR-corrected Wilcoxon Rank-Sum Test p-value < 0.01. b) Quadrant plot exhibiting myeloid cells colored the grades of the associated tumors. c) Boxplot showing the average usage of each program by histopathological tumor grade. d) Boxplot showing module score calculated per tumor in the TCGA LGG-GBM dataset. Score derived using CXCR4 program gene set. e) Boxplot similar to (c) but with cycling program usage. f) Boxplot showing the odds ratio of cycling in each myeloid cell state, calculated independently for each tumor. ‘Cycling’ and program defined by a cell usage >20% of both cycling and indicated program. Increased odds *p<0.05 g) Quadrant plot illustrating cycling cell distribution among programs.

These distinctions in myeloid program composition could be a function of the mutant IDH enzyme or, alternatively, could reflect associations with tumor grade, given that IDH-mutant cohorts include many low grade tumors. In support of the latter, we find that immunomodulatory program composition strongly correlates with grade, with the myeloid composition of Grade 4 IDH-mutant tumors closely approximating Grade 4 IDH-WT tumors (Fig. 3b-c, Extended Data Fig. 6c). Although all IDH-WT gliomas are now designated as grade 4 due to their similarly poor patient outcomes^8^, their histopathological grade was previously incorporated into diagnostic criteria. Examination of a cohort scored with this prior classification revealed that the myeloid program composition of low-grade IDH-WT tumors mirrored that of low-grade IDH-mutant tumors(Fig. 3d). Hence, our data suggest that purported differences in the myeloid compartment of IDH-mutant and IDH-WT are more likely to reflect tumor grade.

Focusing on tumor grade, we found that myeloid cells in high-grade tumors were also enriched for our G2-M cycling program (Fig. 3e). Unexpectedly, a high proportion of these cycling cells expressed the Scavenger Immunosuppressive program (Fig. 3f-g). The Scavenger Immunosuppressive program was the only program enriched for co-expression of cycling programs, whereas the neutrophil and IL1B Inflammatory programs demonstrated minimal overlap with cycling programs.

These results demonstrate that observed differences in the myeloid states in glioma are influenced by grade rather than IDH mutation, and that these differences largely involve differential expression of the immunomodulatory activity programs. This provides a more granular understanding of observations seen in smaller cohorts or with technologies that rely on limited state markers such as multiplex fluorescence and flow cytometry. It also points to the tumor microenvironment as a major driver of the immunomodulatory programs.

### Spatial transcriptomics associates immunomodulatory programs with tumor niches

To investigate potential microenvironmental drivers of the immunomodulatory myeloid programs, we integrated our scRNA-seq data with 10X Visium spatial transcriptomic data^44^. We conducted and combined two independent analyses to relate our cellular programs to tumor niches (Fig. 4a). First, we again leveraged cNMF to identify spatial gene programs in an unbiased manner that were differentially expressed across the 68,830 50 µm pixels in the 23 spatial sections (see Methods). This distinguished six prominent regional expression programs (“niche programs”) that included gray and white matter structural niches, hypoxic and vascular metabolic niches, a niche composed of proliferative cancer cells enriched for genes expressed by OPC-like and NPC-like malignant cells (Proliferative Cancer), and an inflammatory niche composed of immune cells and reactive astrocytic genes (Inflammatory) (Extended Data 7a). In parallel, we estimated the cellular content of each 50 µm pixel by integrating our previously defined scRNA-seq programs (see Methods, Supplemental Table 2) with the spatial data using Robust Cell Type Decomposition (RCTD)^13^. Plotting these data on individual tumor sections revealed clear niche-specific patterns in myeloid programs, cancer cell programs, and other cell types within the tumor (Fig. 4b).

**Fig. 4:**
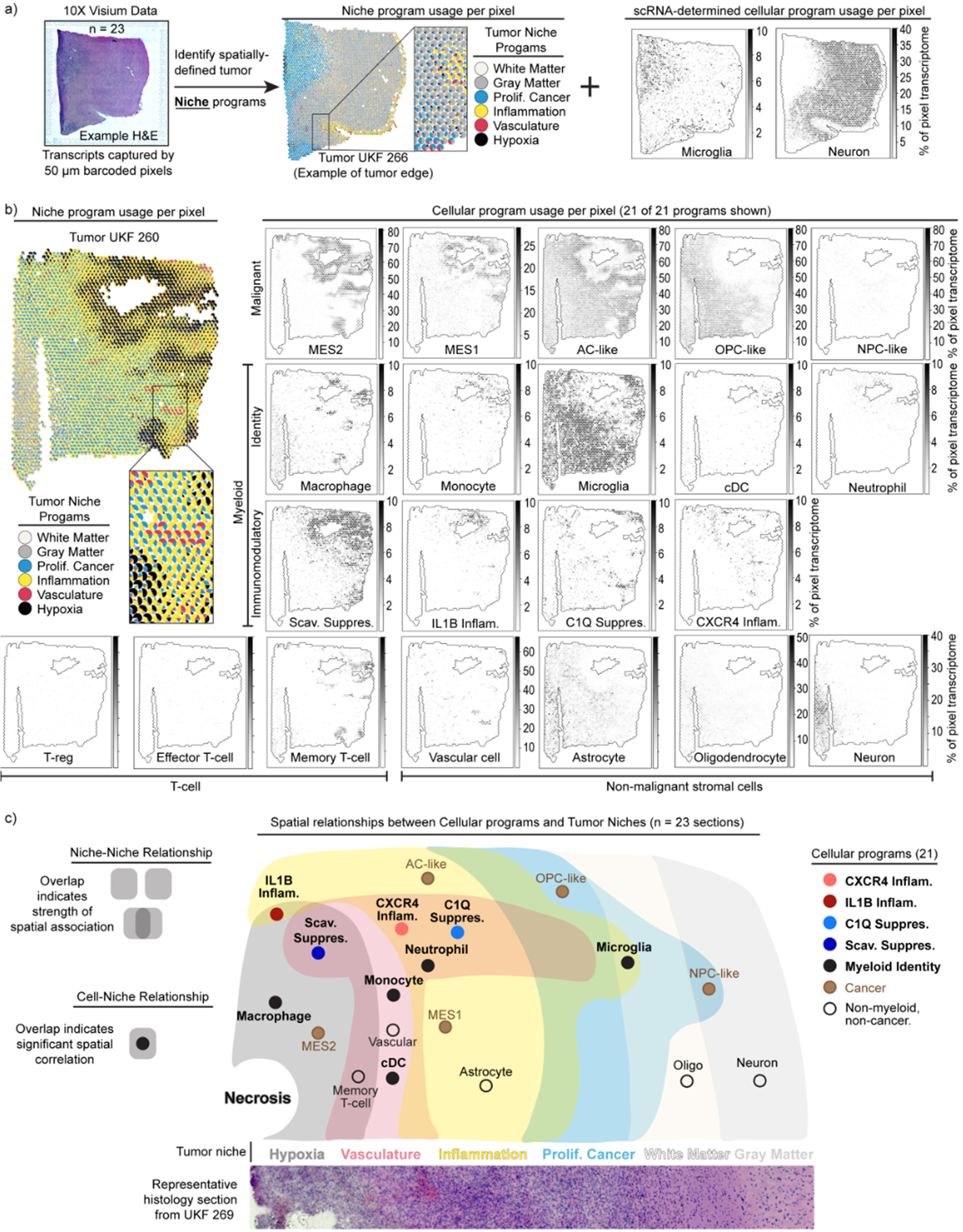
Spatial transcriptomics associates immunomodulatory programs with tumor niches. a) Schematic illustrates the dual analysis approach for spatial transcriptomics samples: cNMF defines broad transcriptomic niches, and RCTD demultiplexes cell content by pixel based on scRNA-seq signatures. The middle and right plots were generated in an identical manner as those in Fig. 4b. b) Scatterpie plot (left) of 10X Visium section. Each pie chart represents a pixel. Scatter plot (right) of the same section. Colors show the RCTD-predicted pixel proportions for adjacent cell types. c) Cell-niche map illustrates conserved spatial relationships of tumor cell types and their ties to transcriptomic niches across spatial transcriptomic samples.

To collate recurrent spatial relationships systematically, we computed intra-pixel correlations between cellular programs and niche programs across all 10X Visium tumor sections. This revealed robust spatial associations between niche programs (niche-niche), between niche and cell programs (niche-cell), and between different cell programs (cell-cell) (Extended Data 7b-d). We derived a single overarching cell-niche map based on the niche-niche and niche-cell associations that showcased these spatial relationships (Fig. 4c).

First, consideration of niche-niche relationships (Extended Data Fig. 7b) reveals a recurrent tumor architecture where a hypoxic niche is flanked by an inflammatory niche, which in turn is adjacent to a proliferative cancer niche that then runs into white matter, consistent with the clinical observation that most gliomas are present in white matter^45^. A vascular niche straddles the hypoxic niche and the inflammatory niche, indicative of vascular proliferation in response to hypoxia and potentially representing an entry point for immune infiltration. These patterns are generally consistent with recent reports^46^.

Second, our assessment of cell-niche relationships (Extended Data Fig. 7c) indicated that the hypoxic regions surrounding necrotic tissue also appear to organize coincident and adjacent cellular programs. Malignant programs were layered around hypoxic regions, with MES2 expressed within the hypoxic niche surrounded by MES1 and AC-like program layers in the Inflammatory niche (Fig. 4b-c). The OPC-like and NPC-like cancer programs were largely excluded from the hypoxic niche and expressed in the proliferative cancer niche.

Hypoxia was similarly organizing for the myeloid cell programs. The Scavenger Immunosuppressive program was almost exclusively found in hypoxic regions, while the C1Q Immunosuppressive program was excluded from hypoxic niches and instead enriched in the surrounding inflammatory and vascular niches (Fig. 4b-c). The IL1B Inflammatory program was associated with hypoxic and Inflammatory niches, while the CXCR4 Inflammatory program was enriched in the Inflammatory and Vascular niches. Microglia were excluded from hypoxic niches, but found throughout the rest of the tumor field. These analyses indicate that each myeloid program has its own tumor niche.

Finally, we used a spatial regression model (see Methods) to assess cell-cell spatial relationships. Our assessment highlighted multiple spatial interactions involving the Scavenger Immunosuppressive program (Extended Data Fig. 7d,e). This program is enriched for spatial interactions with nearly every cell program occupying the hypoxic or vascular niche (Extended Data Fig. 7e). In particular, we noted correlations between the Scavenger Immunosuppressive and the MES2, MES1, and monocyte programs. We validated these connections orthogonally using our complete scRNA-seq dataset, which revealed that average usage of these associated programs was highly correlated with usage of the Scavenger Immunosuppressive program across tumors (Extended Data Fig. 7f). Overall, these data suggest that the Scavenger Immunosuppressive program may be a key determinant of the tumor microenvironment in glioma.

Taken together, our findings propose a consistent and structured tumor architecture across gliomas, with myeloid cell programs demonstrating spatially restricted expression patterns that are associated with and potentially instructed by tumor microenvironmental cues. In particular, metabolic factors (e.g., hypoxia, vascular), proximal cell states (e.g., MES2), and brain structure (e.g., gray matter, white matter) appear to direct alternate myeloid programs. This raises the interesting corollary that the extent of tumor resection dictates which microenvironments and associated myeloid cell programs remain following surgery, and that incomplete resection of hypoxic regions results in increased presence of the Scavenger Immunosuppressive program.

### Dexamethasone drives the C1Q Immunosuppressive program

In addition to tumor niches, we asked if clinical therapies might have an effect on myeloid cell states. Dexamethasone is a potent corticosteroid routinely administered to glioma patients to reduce tumor-induced vasogenic edema in the brain pre- and post-operatively. Given that dexamethasone is also used to suppress inflammation in many diseases, we postulated it may be influencing myeloid cell programs. Therefore, we first tested if the dose of dexamethasone was significantly associated with any of our myeloid programs. In the MGB and McGill cohorts where treatment information was accessible to us, we find that the C1Q Immunosuppressive program is specifically and significantly associated with increasing steroid dose (Fig. 5a-b). Subsequently, we leveraged our MGB cohort dataset to contrast the myeloid profiles of patients treated with and without dexamethasone. This unique cohort included multiple patients who were not treated with dexamethasone due to concerns that the agent might hinder response to post operative immunotherapy trials. We find a specific and statistically significant association between use of dexamethasone and the C1Q Immunosuppressive program when controlling for the confounding effect of hypoxia (Fig. 5c).

**Fig. 5:**
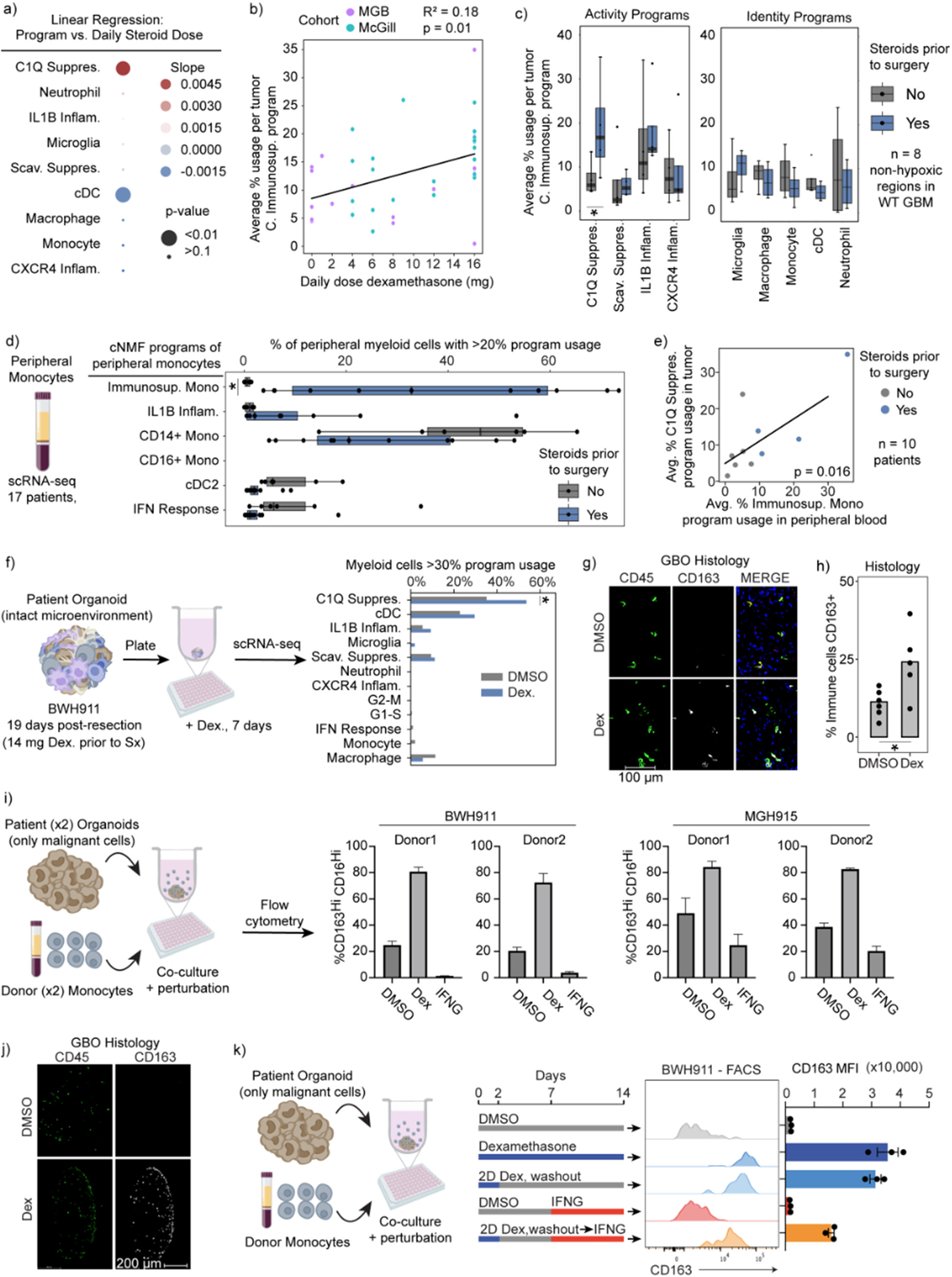
Dexamethasone drives the C1Q Immunosuppressive program. a) Dot plot displays the linear regression coefficient between each myeloid program’s average usage per sample and the respective patient’s pre-surgery daily dexamethasone dose, using only IDH-WT samples. b) Scatterplot of mean C1Q Immunosuppressive program with least-square linear regression line. c) Boxplot displays the average usage of programs stratified by use of dexamethasone in IDH-WT tumors with low hypoxic program usage in the MGB cohort. d) Boxplot of the percent of myeloid cells with the indicated peripheral myeloid programs in peripheral myeloid cells from patients with gliomas. e) Scatterplot illustrates the average C1Q Immunosuppressive usage in myeloid cells of tumor samples versus average Immunosuppressive Monocyte usage in related peripheral myeloid cells. Only tumors with low hypoxic program usage are considered. f) Schematic (left) and bar graph (right) of the percentage of myeloid cells expressing the C1Q Immunosuppressive program. P-value obtained using Fisher’s Exact test. * p-value <0.05, all others have p-value > 0.2. g) Immunofluorescence image of a GBO with intact endogenous TME co-cultured for 7 days with DMSO or 100 nM dexamethasone. h) Quantification of marker positive cells in sectioned organoids. Each dot represents an organoid in the condition. Student’s T-test p<0.05. i) Schematic (left) and bar plot of flow cytometry results from experiment. Error bars St. Dev. j) Representative section of organoid and infiltrated monocytes when treated with dexamethasone. k) (left) Schematic of experimental design. (right) Flow cytometry results. Error bars St.Dev. Unless otherwise indicated *FDR-corrected Wilcoxon Rank-Sum Test p-value < 0.05.

Given that both dexamethasone and myeloid cells can originate from blood, we next investigated whether dexamethasone also triggers suppressive phenotypes in circulating monocytes. We turned to our scRNA-seq data of peripheral blood of glioma patients. Stratifying patients by dexamethasone treatment, we again found one program in peripheral monocytes specifically increased in patients treated with dexamethasone (Fig. 5d). This program includes *CD163* and other markers found in the C1Q Immunosuppressive program, although it was not completely overlapping, raising the possibility that this is a precursor program in the periphery to the program that develops in myeloid cells in the tumor. We also observed a positive correlation between the average expression of the dexamethasone-related program in circulating monocytes and the average expression of the C1Q Immunosuppressive program in tumor-associated monocytes from the same patient (Fig. 5e).

To test whether dexamethasone can directly drive expression of the C1Q Immunosuppressive program in myeloid cells, we turned to our tumor organoid systems. We focused initially on endogenous tumor myeloid cells in organoids from recently resected tumors that maintained the original tumor microenvironment, including myeloid cells. Dexamethasone specifically induced the C1Q Immunosuppressive program per scRNA-seq (Fig. 5f). CD163, a surface protein marker associated with both of our immunosuppressive programs, was also increased (Fig. 5g,h). We also modeled infiltration of peripheral myeloid cells into the tumor by adding peripheral human monocytes to tumor organoids devoid of immune cells. Dexamethasone again strongly induced expression of the C1Q Immunosuppressive program in myeloid cells that infiltrated into the organoid (Fig. 5i,j).

Mirroring the real-world scenario where patients receive corticosteroids preoperatively, only to be discontinued post-surgery, we investigated whether dexamethasone-induced changes were reversible. We treated myeloid cells infiltrating tumor organoids for 2 days with dexamethasone and then washed out the drug from the wells and waited 2 weeks. Importantly, we found C1Q immunosuppressive program expression did not reverse even 2 weeks after drug withdrawal (Fig. 5k and Extended Data Fig. 8). This dexamethasone-induced state change was only partially rescued by addition of high level IFN-γ.

Altogether, these data indicate that dexamethasone drives the C1Q Immunosuppressive program in gliomas in a largely irreversible manner and may also create a pool of circulating suppressive monocytes that subsequently infiltrate tumor.

### Clinical correlates with immune suppression and patient outcomes

Finally, we sought to relate our glioma-associated myeloid programs to clinical correlates of immunity and outcome. Focusing first on infiltrating T cells (see Methods, Supplemental Table 2), we found that a majority expressed signatures consistent with Naive/Memory T-cells (65%), while 25% resembled Effector T-cells and another 7% T regulatory cells (T-reg) (Extended Data Fig. 9a). We did not detect a prominent program for exhausted T-cells, suggesting that this population is rare in our cohort. Given their established links to myeloid cells and immunosuppression^47–49^, we related T-reg proportions to our programs. Tumors with high T-reg frequency were enriched for myeloid cells expressing Scavenger and C1Q Immunosuppressive programs, but were depleted of CXCR4 Inflammatory-expressing cells (Fig. 6a). We also detected a spatial association between T-reg and the C1Q Immunosuppressive program (Extended Data Fig. 7d-e), suggesting that T-reg cells reside in close proximity to C1Q-expressing myeloid cells. In contrast, T-cells with Naive/Memory expression signatures were spatially associated with hypoxic niches and the Scavenger Immunosuppressive program (Fig. 4c, Extended Data Fig. 7c). These results suggest that the respective immunosuppressive myeloid programs distinctly impact T-cell states and the immune microenvironment in gliomas.

**Fig. 6:**
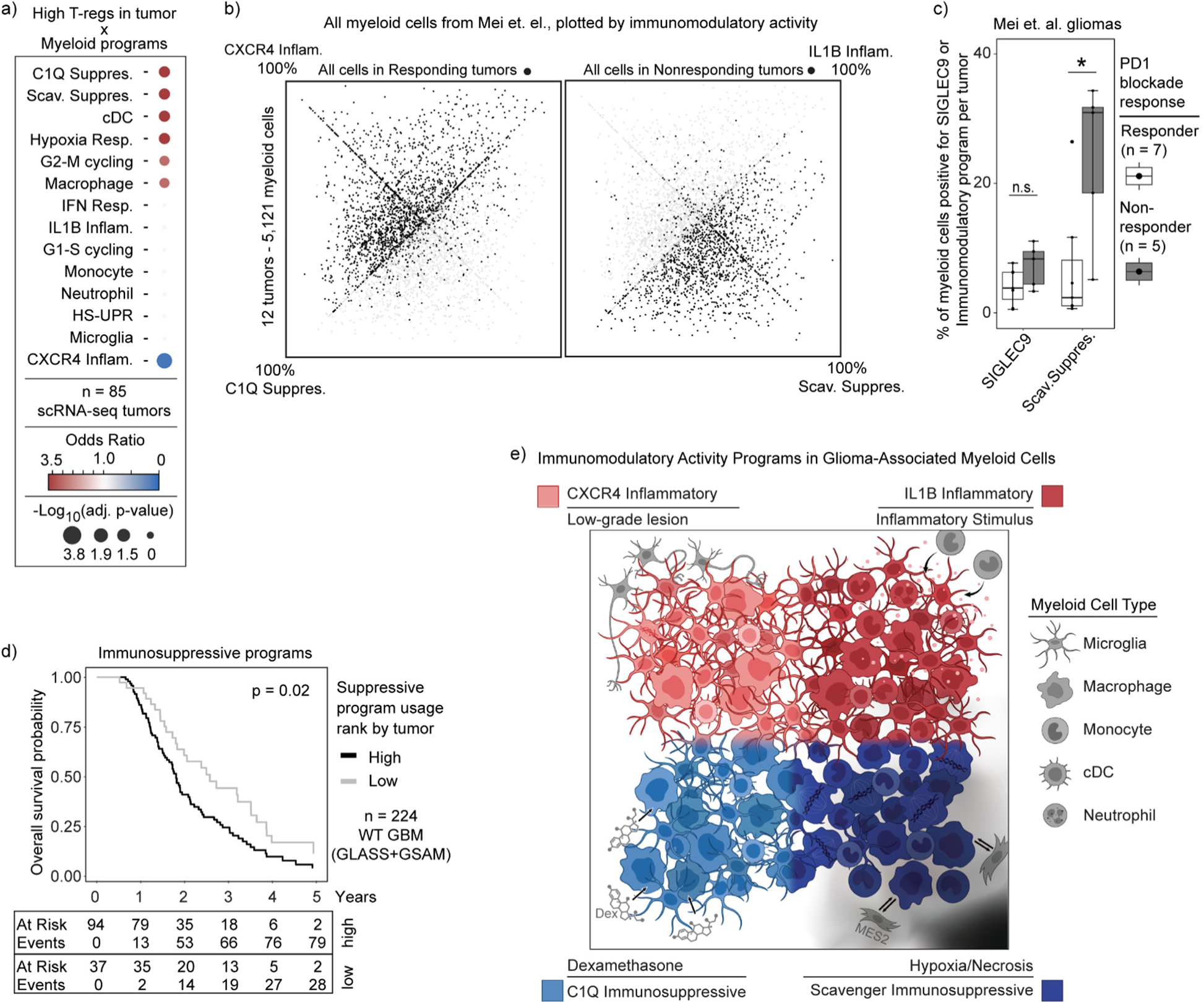
Immunosuppressive programs associated with immunotherapy resistance and worse overall survival. a) Dot plot displaying the odds ratio for high program expression in tumors with high T-reg abundance. b) Quadrant plot of cells from Mei et. al., plotted based on expression of our immunomodulatory activity programs, highlighting cells in tumors with response or nonresponse to immunotherapy. c) Boxplot of per tumor calculation of SIGLEC9-positive cells or Scavenger Immunosuppressive program usage > 20%. d) Kaplan-Meyer curve of overall survival by combined immunosuppressive program expression. P-value calculated using the Cox proportional hazards regression model. e) Summary figure.

We next investigated whether our myeloid programs predict response to immunotherapy. A recent scRNA-seq study of 12 glioma patients treated with neoadjuvant PD1 blockade identified a population of SIGLEC9-expressing macrophages that accumulated in non-responsive tumors^50^. Reanalysis of these data using our cNMF framework revealed that SIGLEC9 positive cells were heterogeneous in their expression of our immunomodulatory programs (Extended Data Fig. 9b). Notably, only the SIGLEC9-positive cells expressing the Scavenger Immunosuppressive program were enriched in non-responders (Extended Data Fig. 9c), and that usage of the Scavenger Immunosuppressive program on its own was more closely associated with non-responding gliomas than SIGLEC9 alone (Fig. 6b-c, Extended Data 9b,d), indicating this immunosuppressive program may more fully explain the immunotherapy resistance phenotype than SIGLEC9 alone. Indeed, there was a striking difference in overall distribution of our immunomodulatory program usage in cells in responsive versus non-responsive tumors (Fig. 6b). This analysis highlights a potentially critical role for this myeloid program in suppressing T-cell activation and/or other key determinants of response to checkpoint therapy.

Finally, we asked whether any of our four myeloid cell programs were associated with survival. We used the top genes of our myeloid programs to score each tumor and adjusted the results based on its estimated myeloid content (see Methods). To avoid confounding effects of tumor grade and IDH mutation status, we limited our analysis to IDH WT glioblastoma patients. We found that high expression of the C1Q and Scavenger Immunosuppressive programs was significantly associated with worse overall patient survival (Fig. 6d), whereas no other myeloid programs were significant, suggesting that immunosuppressive myeloid microenvironments may be detrimental to survival even in the absence of immunotherapy.

In summary, our analysis of clinical specimens and correlates suggests that the C1Q and Scavenger Immunosuppressive myeloid programs may shape T-cell phenotypes in the tumor microenvironment and, moreover, impact patient outcome and response to immunotherapy. While we cannot rule out that the associations may be partially correlative, prior literature and our spatial findings support causal roles for the myeloid programs in shaping the glioma microenvironment and these functional outcomes.

## DISCUSSION

Harnessing the power of the immune system is arguably the most promising path to a cure for glioma patients. However, therapeutic development has been hindered by the unique immune microenvironment of brain tumors, which are densely infiltrated with myeloid cells and depleted of T cells. Here we combined single-cell and spatial genomic technologies for more than 100 tumors, a complementary computational framework, clinical data, and functional experimental models to create foundational insights into myeloid cells in glioma. We detail the spectrum of glioma-associated myeloid cell types, their developmental origins, and the immunomodulatory programs that they express. It answers key biological questions and should catalyze basic and translational efforts going forward.

Our study highlights the plasticity of glioma-associated myeloid cells and the impact of the local microenvironment on their phenotypes. By decomposing scRNA-seq data of each cell into unbiased, discrete gene expression programs using cNMF, we disentangle cell identity from cell activity. This change in approach unveiled previously obscured biological insights. Although myeloid cells have typically been classified by cell type or cell ontogeny, we found neither to be major determinants of myeloid cell activity in gliomas. Rather, different myeloid cell types, including microglia, macrophages, and monocytes, can each engage the same set of immunomodulatory activity programs. Activation of each of these four programs appears to be largely determined by unique drivers in the microenvironment (Fig. 6e). The immunosuppressive programs are independently associated with either hypoxic regions in the tumors (Scavenger) or dexamethasone treatment (C1Q). The CXCR4 Inflammatory program is associated with low grade lesions where interactions with non-malignant neural cell types are prevalent, while the IL1B Inflammatory program appears to be a default program in response to an inflammatory microenvironment and itself seems to recruit additional myeloid cells into the tumor.

Further indication of myeloid cell plasticity emerged from our inferential analysis of developmental origins on the basis of mitochondrial DNA mutations. This analysis revealed that blood-derived monocytes can adopt microglia-like expression states in tumors and that both blood-derived and resident cells can activate the full range of immunomodulatory programs. Moreover, we found that peripheral blood monocytes can rapidly differentiate and activate the different immunomodulatory programs when applied to glioma organoids. These findings underscore the potency of the tumor microenvironment for programming the functional phenotypes of myeloid cells, and stress the need for caution when inferring cellular origin from markers or immune function on the basis of myeloid cell type. They provide incentive to develop an updated model of myeloid cell development and phenotypes in the injured human brain.

Based on our findings, we propose the following framework for glioma-associated myeloid cells, which may also be applicable to brain metastases and potentially other cancer types. First, myeloid states are composed of superimposable identity and activity programs and should be characterized and annotated accordingly. Second, myeloid cells exhibit striking developmental and phenotypic plasticity. Tumor niches potently influence their differentiation trajectories and immunomodulatory programs. Third, myeloid immunomodulatory programs shape the overall immune state of gliomas and, as such, are associated with patient outcome and response to immunotherapy. Fourth, the immunomodulatory programs and potentially the underlying cell states can be modulated by clinical and experimental interventions. Finally, our analyses suggest that therapeutic interventions should target specific immunomodulatory programs rather than indiscriminate myeloid cell targeting. Accordingly, our framework for systematic annotation and characterization of myeloid states in tumors and experimental models can catalyze and harmonize the study of myeloid programs and interventions, including studies that aim to modulate the immune microenvironment for therapeutic gains. In addition to its critical mass of data and program definitions, our resource includes a cloud-based pipeline and portal for exploration of our data, and for the integration and analysis of additional datasets within this framework.

This framework complements and builds on prior studies that provided evidence for the diversity and plasticity of glioma-associated myeloid cell states^4,7,18–21,23–25^, and hinted at the convergence of myeloid programs in response to the microenvironment^18,51^. In particular, a bulk RNA-seq analysis of sorted cell populations by Klemm and colleagues^18^ revealed that microglia acquired monocyte-derived macrophage features in IDH WT tumors and brain metastases, while monocyte-derived macrophages acquired microglia features in IDH-mutant gliomas. The importance of microenvironment was also highlighted in seminal work finding that resident myeloid phenotypes in non-cancerous tissue are shaped by their local microenvironment more than origin^51^. However, the field has been slowed by an inability to effectively disentangle cell type from activity and by limited sample sizes, with prior studies coming to different conclusions in several areas. Multiple studies have concluded that IDH mutation directly drives differences in myeloid cell phenotype^4,18,21^, while our analysis suggests this is largely driven by the microenvironment associated with tumor grade. Other studies found differences in myeloid phenotype with recurrence^23^, which we do not see in our larger cohort. Finally, while most studies focus on origin or cell type as the distinguishing features of myeloid cell states^18,23–25^, we show that specific immunomodulatory programs are shared across different myeloid cell types in response to the tumor microenvironment, regardless of origin. This convergence of immune phenotypes is highly significant for glioma biology and treatment.

We recognize that our study has significant limitations and leaves critical questions unaddressed. Our data capture the diversity of myeloid cells at the time of tumor resection, but cannot appreciate their temporal dynamics, the stability of the myeloid cell programs in a given cell, or the rate of myeloid cell turnover in glioma. These fundamental questions regarding myeloid cell plasticity are clinically important and particularly timely given recent mouse modeling studies suggesting that myeloid cell lifespan may be extended in brain tumor niches^52^. Further study is also needed to define the specific signaling molecules that drive myeloid cell invasion, differentiation, and immunomodulatory program usage in gliomas. Although our study hints at the potential of therapeutic interventions to modulate glioma-associated myeloid states and the broader immune environment, their rational development will require insights into these signaling mechanisms and the downstream transcriptional and epigenetic regulators that create and maintain the immunomodulatory programs. A clearer understanding of these myeloid programs and determinants will require definition and integration of astrocytes, endothelial cells, pericytes and other immune and non-immune cell types in the glioma ecosystem^53^.

In conclusion, we highlight potential clinical implications. Most addressable is the C1Q Immunosuppressive program, which is irreversibly driven by dexamethasone. This effect likely impacts immunotherapy clinical trials, as most permit some level of dexamethasone use. It also highlights the importance of creating alternatives to dexamethasone for symptom management. Finally, our data nominates the Scavenger Immunosuppressive program as a target for future work given its associations with immunotherapy resistance and poor outcomes. We hope that these foundational datasets and framework can harmonize and catalyze the study of brain tumor myeloid cells and pave the way for therapeutic strategies designed to alter tumor microenvironments to increase immunotherapy efficacy.

## Extended Data

**Extended Data Fig. 1:**
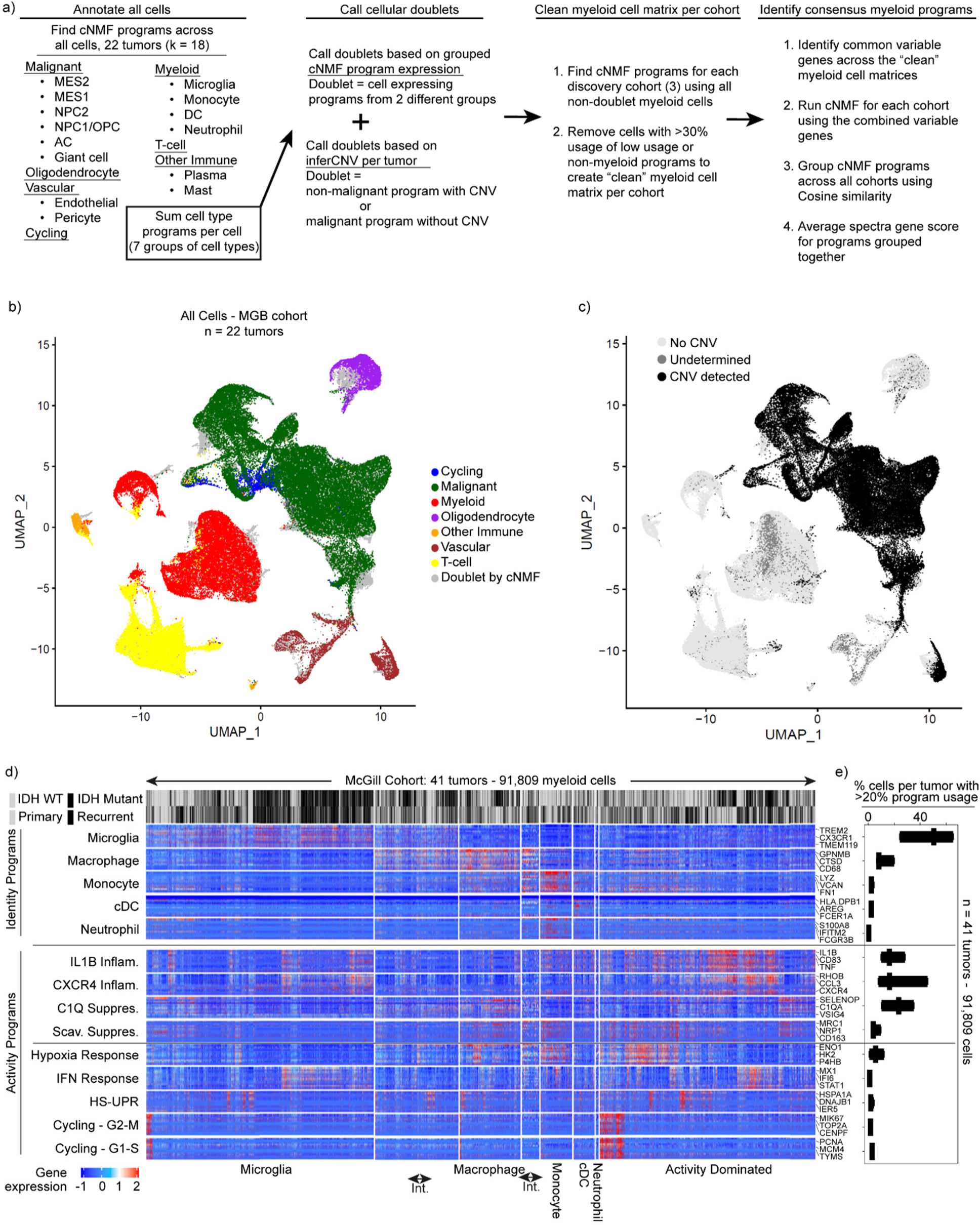
Identification of consensus myeloid programs and validation. a) Schematics of the computational pipeline for the identification of the recurrent myeloid programs across the scRNA-Seq libraries of the three discovery cohorts. b) UMAP of the broad annotation of all cells in the MGB cohort according to the key. c) UMAP demonstrating the presence (black) or absence (gray) of copy number variation events in all cells of the MGB cohort tumors. d) Heatmaps demonstrating the expression of genes in recurrent myeloid programs (rows) by cell (column) grouped by cell type. Cell usage of myeloid identity programs was used to define myeloid cell type e) Boxplots exhibiting the percentage of cells by sample expressing the recurrent myeloid programs indicated on the left of the heatmap across the validation cohort.

**Extended Data Fig. 2:**
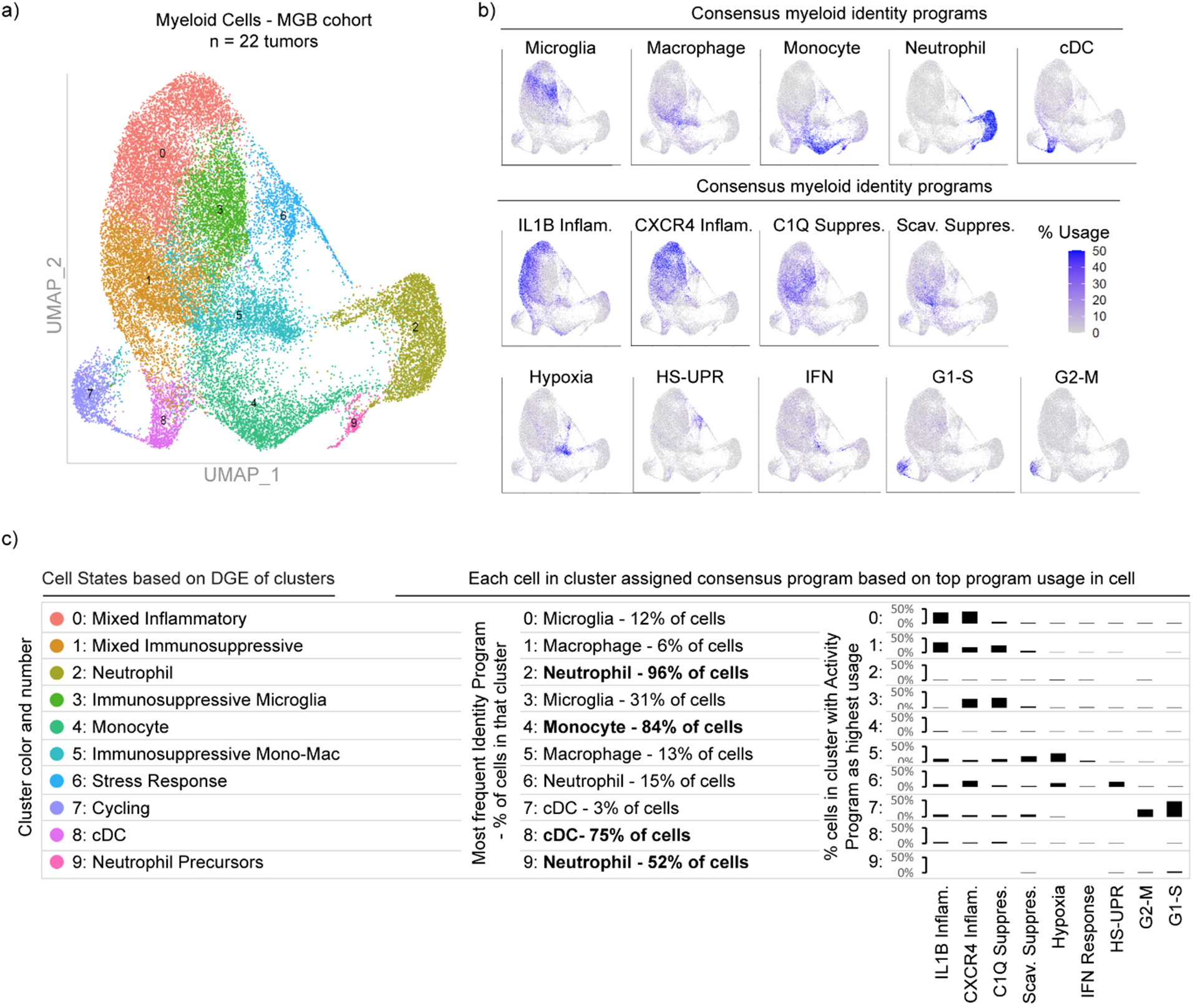
Direct comparison of Louvain clustering and cNMF programs. a) UMAP exhibiting the Louvain clusters of batch-corrected singlet myeloid cells of the MGB cohort. b) UMAPs of the myeloid cells of the MGB cohort demonstrating the usage of indicated programs at the top of each UMAP. c) (left) Annotations of Louvain clusters in (a) based on standard differential gene expression analysis of clusters. (Center) Name and frequency of most frequent cell type in the Louvain cluster as annotated by cNMF identity programs. (Right) bar chart of the percent of cells in the Louvain cluster with a given activity program as top program used in that cell, based on cNMF programs.

**Extended Data Fig. 3:**
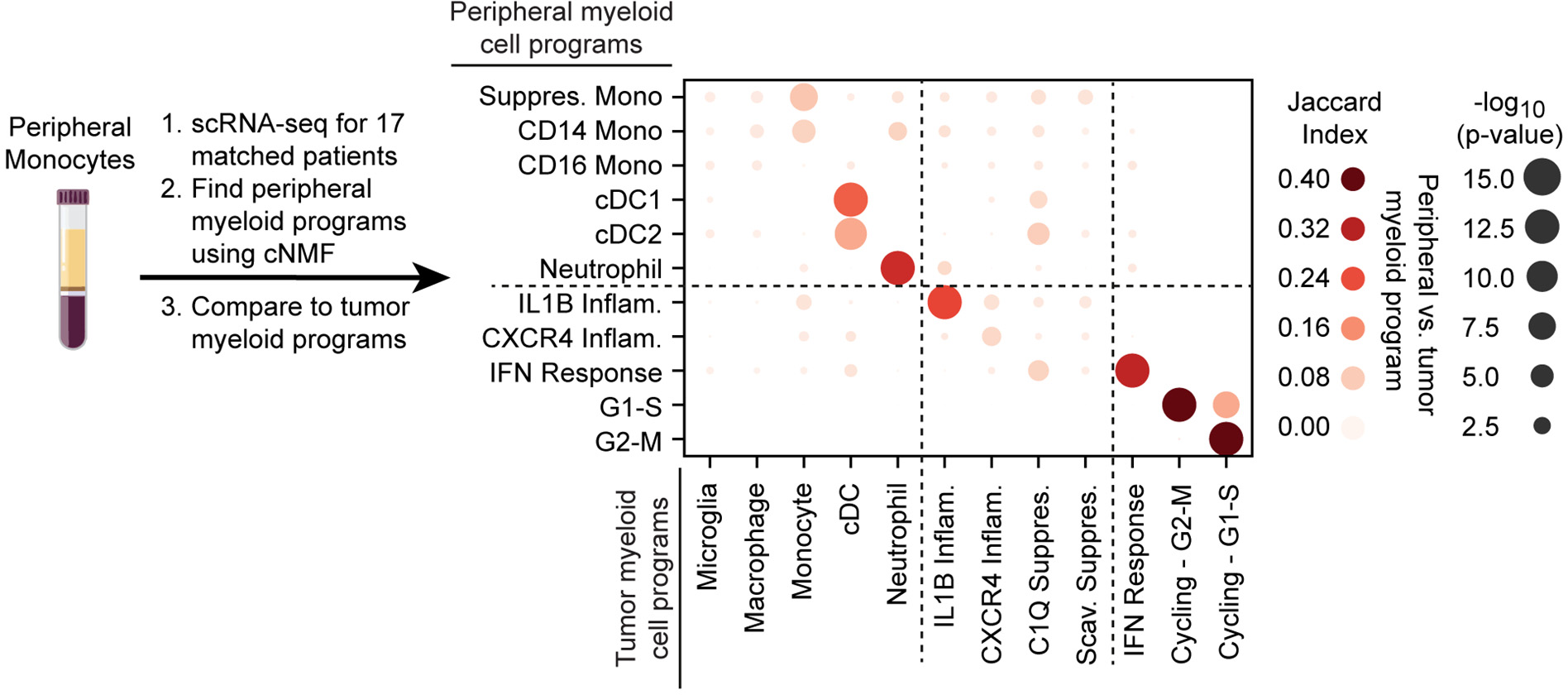
Identification and comparison of peripheral myeloid cNMF program to tumor cNMF programs. (Left) Schematic of peripheral myeloid cell program identification. (Right) Dot plot of Jaccard Index between peripheral myeloid cNMF programs and tumor cNMF programs.

**Extended Data Fig. 4:**
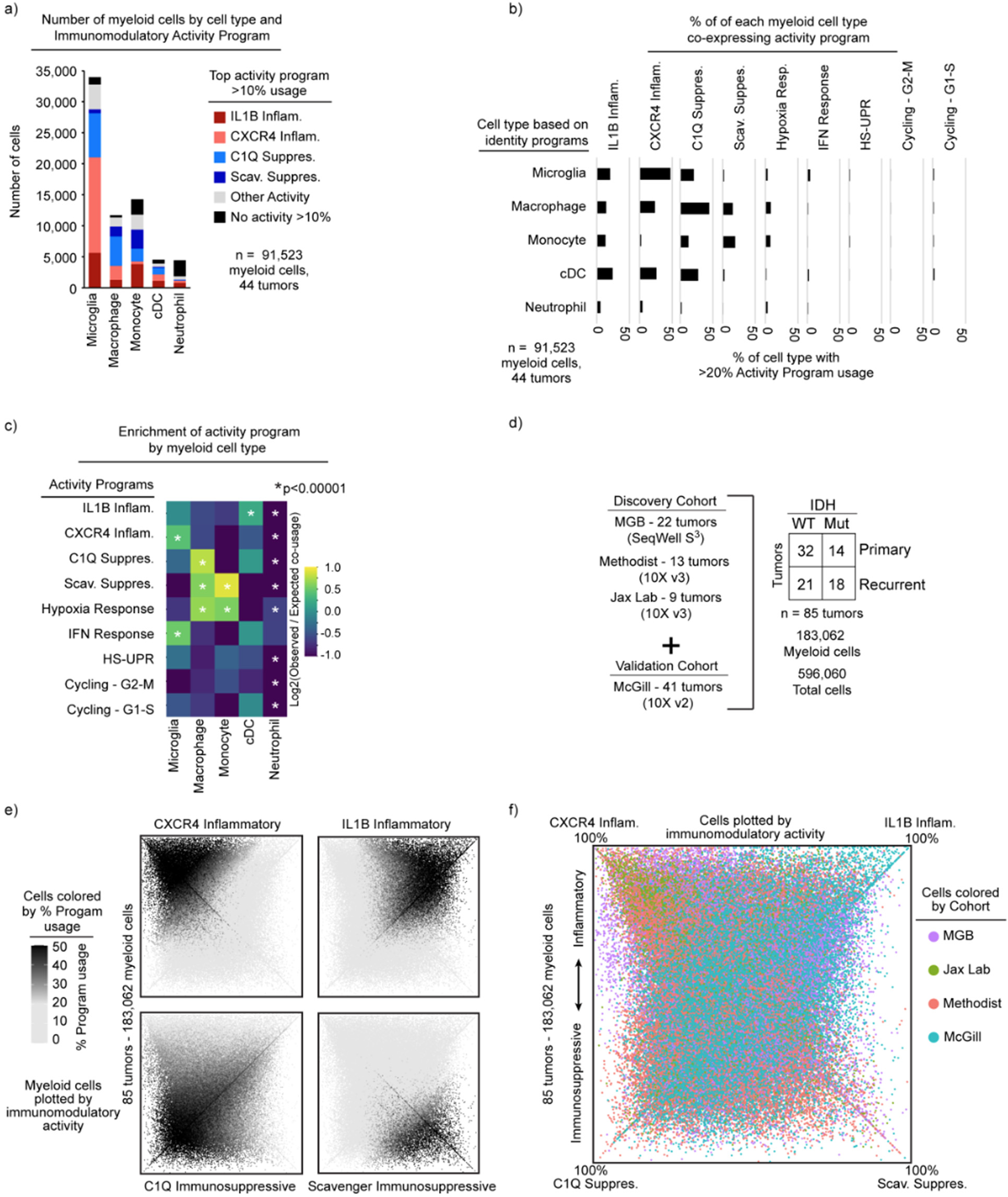
Activity program usage among myeloid cell types. a) Stacked bar plots of absolute number of cells with activity program usage per myeloid cell type. b) Horizontal bar chart of percent of cell type with >20% activity program usage. c) Enrichment plot demonstrating the enrichment level of the four immunomodulatory programs (left) in the shown identities (above). d) Schematics demonstrating the inclusion of the McGill Validation cohort in all subsequent analyses. e) Quadrant plots in which the color represents the usage level of the indicated immunomodulatory program. f) Quadrant plot in which the color represents the cohort from which the myeloid cell comes. The position of each dot represents the difference in the usage of immunosuppressive and inflammatory programs by that cell (the upper part of the plot is more inflammatory, vs. the lower part is more immunosuppressive).

**Extended Data Fig. 5:**
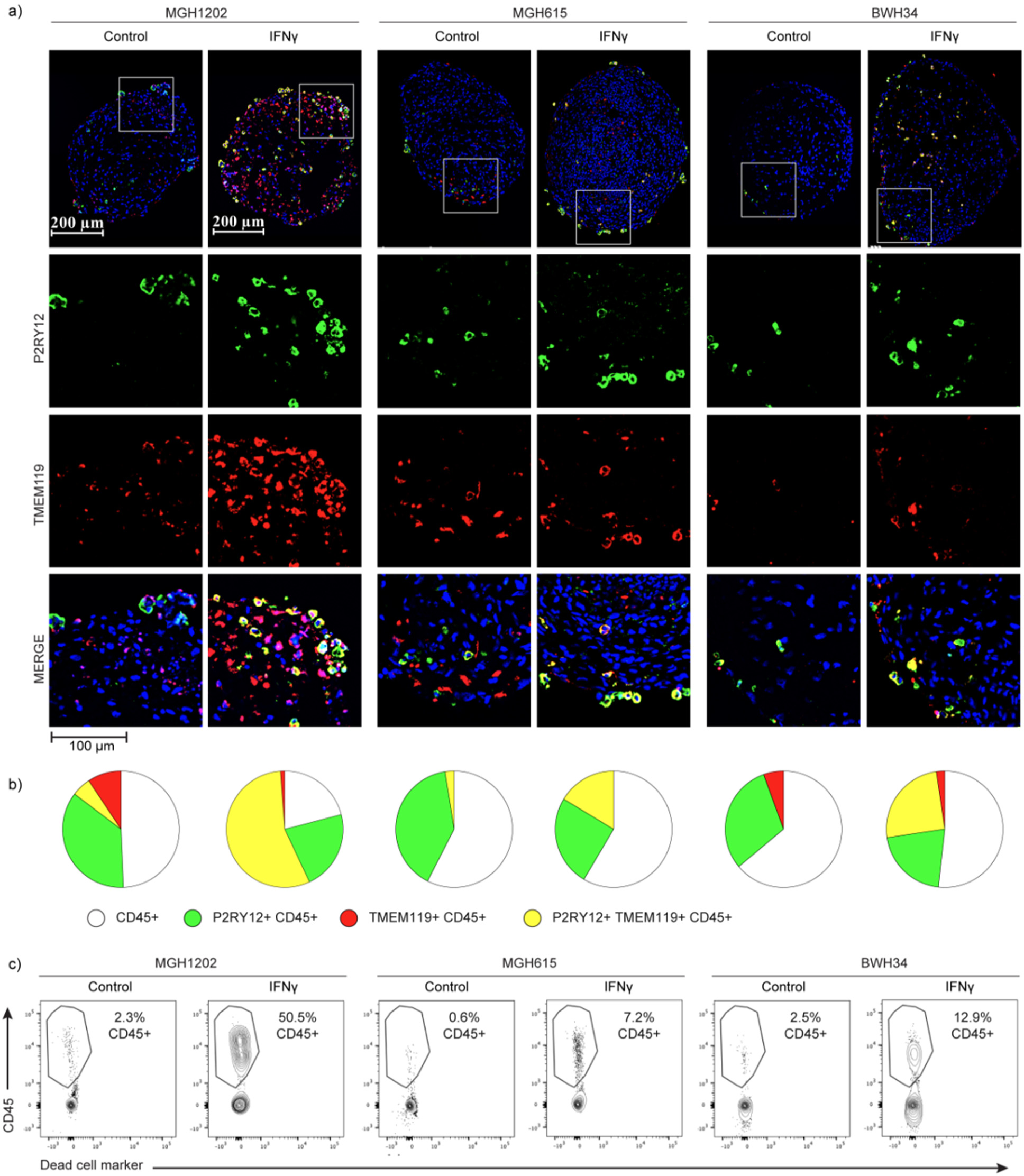
Peripheral monocytes differentiate to express microglia markers in tumor microenvironment, which is potentiated by interferon. a) Representative immunofluorescence images of organoid sections from experiments related to Fig. 2d,e. b) Quantification of images in (a). c) Flow cytometry results of percent of CD45+ cells infiltrating into the organoids. Results are from multiple organoids mixed together for each condition.

**Extended Data Fig 6:**
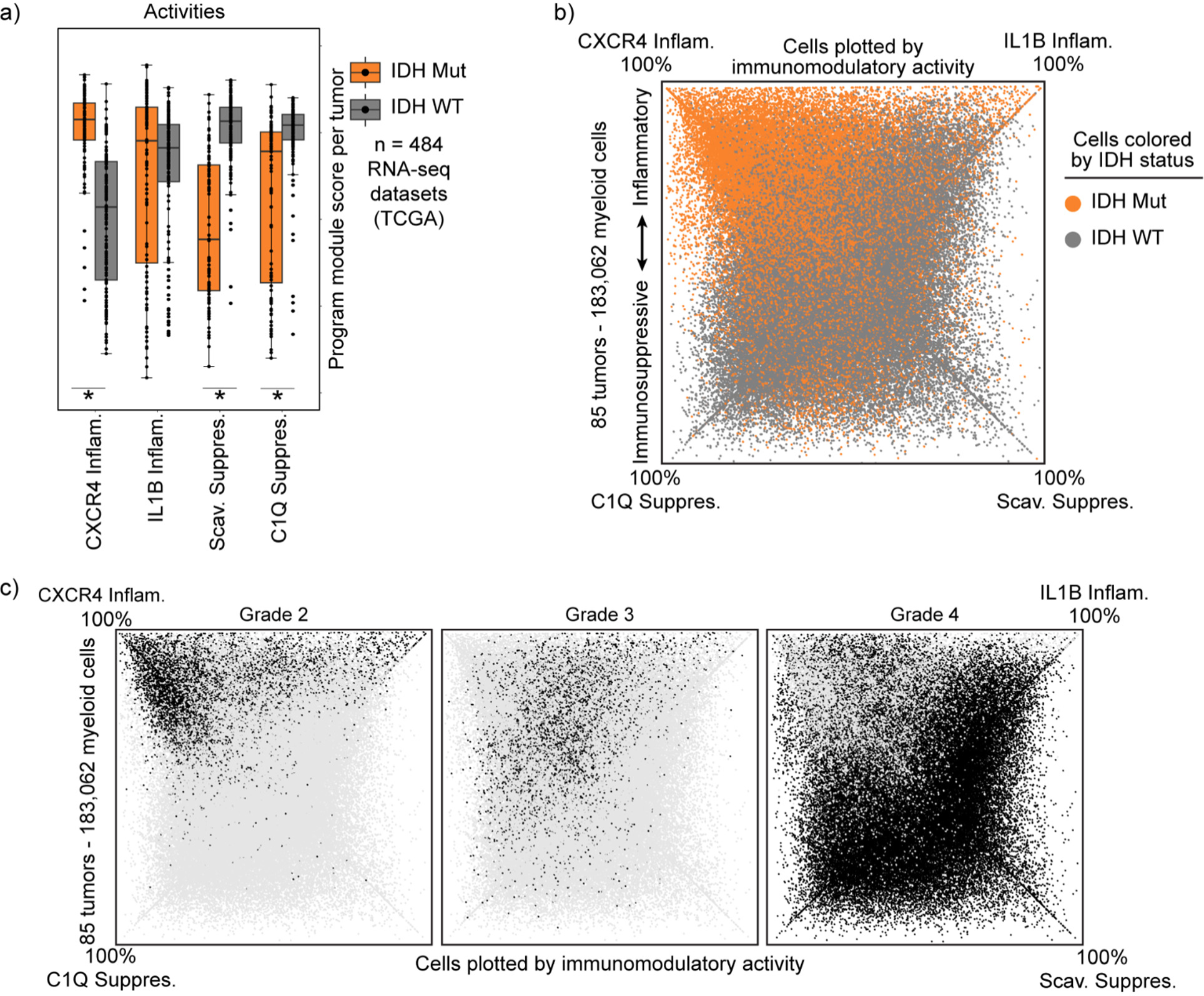
Program composition varies with histopathological tumor grade more than IDH-mutation status. a) Box plot exhibiting the program module scores in tumors of the TCGA cohort. Each dot represents a tumor. * FDR-corrected Wilcoxon Rank-Sum Test p-value < 0.05. b) Quadrant plot exhibiting the IDH mutation status of the tumors from which the myeloid cells come. c) Quadrant plots exhibiting the grade of the tumors from which the myeloid cells come. Black denotes that the myeloid cell comes from a tumor with the grade displayed at the top of each quadrant plot, whereas grey indicates not that grade.

**Extended Data Fig. 7:**
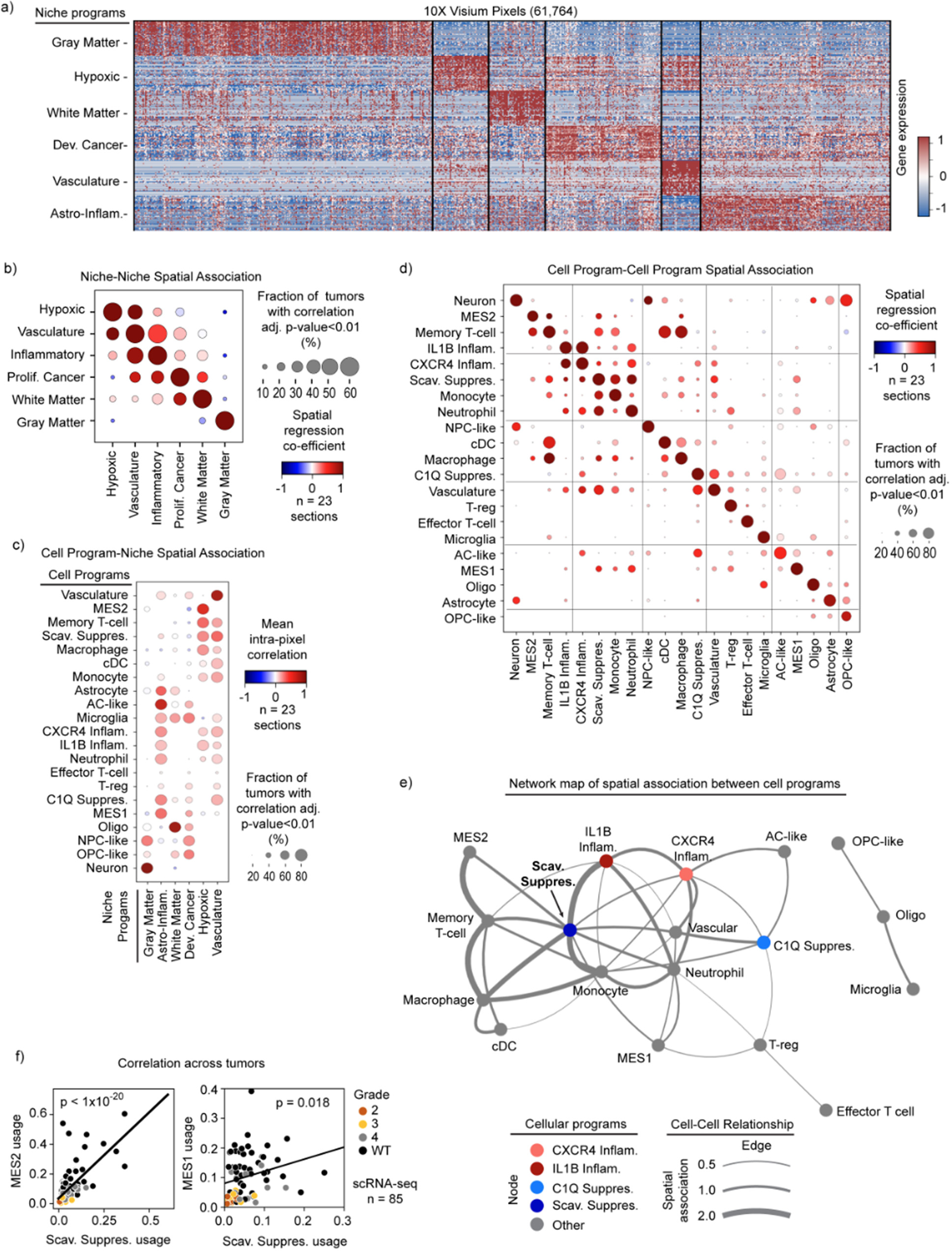
Spatial associations of cells and niches in glioma. a) Heatmap shows gene (rows) expression across all pixels (columns) in the cohort of spatial transcriptomic samples. Top 40 genes of each niche program are shown. Gene expression data is cell normalized, then log normalized and scaled by variance. b) Dot plot displays the spatial proximity enrichment score between niche programs, calculated independently per sample (see Methods for details). Dot size denotes the proportion of samples showing a significant correlation (p-adj < 0.01), while color signifies a positive (red) or negative (blue) correlation. c) Dot plot represents intra-pixel correlation between niche and cell type scores, calculated independently for each sample. Dot size shows the proportion of samples with a statistically significant correlation (p-adj<0.01), while color indicates a positive (red) or negative (blue) correlation. d) Dot plot displays the spatial proximity enrichment score between cell programs, calculated independently per sample. e) Network graph illustrates recurrent spatial relationships of tumor cell types across spatial transcriptomic samples. Nodes denote cell types, with edges marking significantly enriched proximities between cell types, observed in at least 40% of samples with an average enrichment score of at least 0.1. Edge width reflects this average score. f) Scatter plot exhibits the mean Scavenger Immunosuppressive program score (x-axis) versus the MES2 or MES1 cancer program score (y-axis) in the scRNA-seq dataset. Linear least square results are shown (line, and p-value).

**Extended Data Fig. 8:**
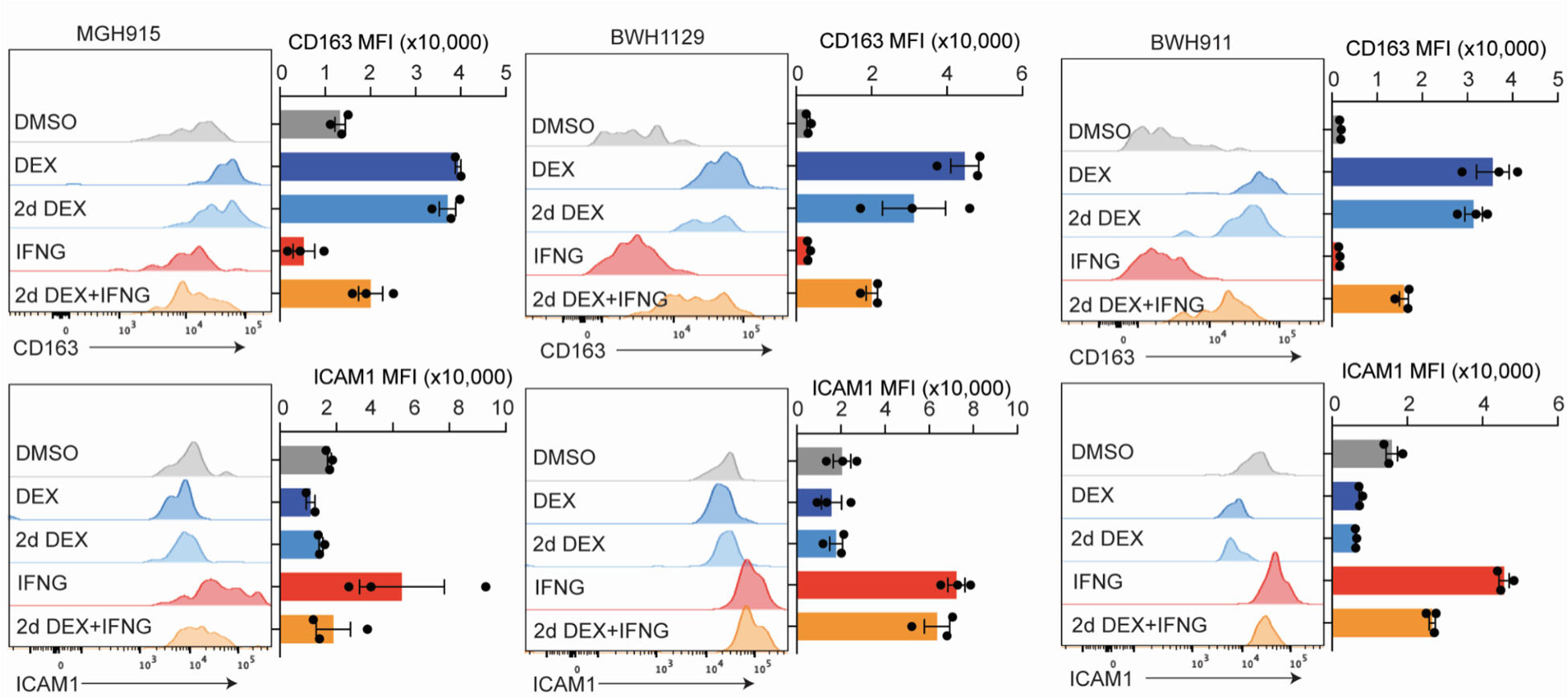
Additional patient organoid models show irreversible phenotype. Flow cytometry results with organoids from multiple patients show the same phenotype as Fig. 5k. Bottom row shows ICAM1, a marker of the IL1B Inflammatory program.

**Extended Data Fig. 9:**
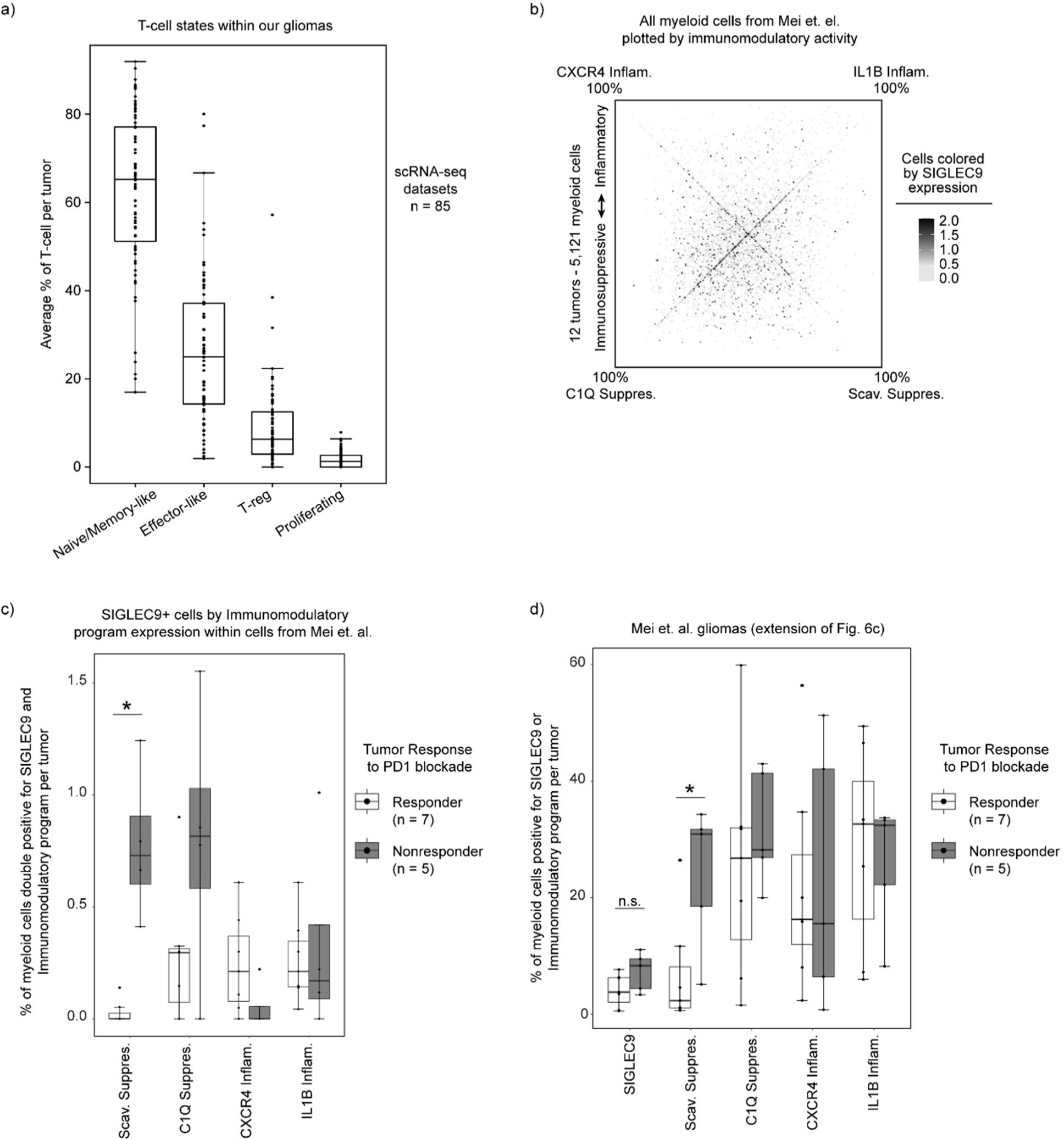
Comparison of SIGLEC9 and Immunomodulatory programs in relation to immunotherapy resistance. a) Boxplot of percent of T cell state per tumor from our scRNA-seq datasets. b) Quadrant plot of cells from Mei et. al., plotted based on expression of our immunomodulatory activity programs, highlighting SIGLEC9 expression heterogeneity. c) Boxplot of SIGLEC9-positive cells from Mei et. al. that were grouped by expression of our immunomodulatory programs, then divided by corresponding tumor response to immunotherapy. Average per tumor plotted. d) Boxplot of per tumor calculation of SIGLEC9-positive cells or Scavenger Immunosuppressive program usage > 20%.

## Acknowledgements

The authors thank Dr. Bo Li for technical help with our Cumulus pipeline and Dr. Itay Tirosh and all Bernstein lab members for critical feedback on the project. We thank patients and their families for generously donating tumor tissue for this study. We thank Julia M. Larson, Magali A. De Sauvage, Elizabeth Summers, the MGH Pathology Tissue Bank, and MGH Neurosurgery, along with Michael C. Prabhu, Connor C. Bossi, and Dr. Keith L. Ligon, and Neurooncology Tissue and Data Bank for their help acquiring fresh tissue resections.

T.E.M. and this work were supported by the UK Brain Tumour Charities Future Leaders Award, GN-000701, the American Brain Tumor Association Basic Research Fellowship in honor of Joel A. Gingras, Jr., the NCI K08 Award K08CA276819, and the NIH T32 Training Grant CA9216. C.P.C. was supported by postdoctoral fellowships from the Ludwig Center at MIT’s Koch Institute, the Canadian Institutes of Health Research (181907), and the Fond de Recherche du Québec - Santé, as well as a grant from the J.H. and E.V Wade Fund at MIT. Z.C. was supported by NCI-CA-234842. D.S.F. was supported by the Eric and Wendy Schmidt Center at the Broad Institute of MIT and Harvard. J.L.G. is supported by R37CA269499.. This project was supported by funds from The Brilliant Night Foundation (K.P.), the NCI/NIH Director’s Fund (DP1CA216873 to B.E.B.), the Ludwig Center at Harvard, and the Emerson Collective. B.E.B. is the Richard and Nancy Lubin Family Endowed Chair at the Dana Farber Cancer Institute, an American Cancer Society Research Professor, and an Investigator in the Ludwig Center at Harvard

## METHODS

### LEAD CONTACT AND MATERIALS AVAILABILITY

Further information and request for resources and reagents should be directed to Bradley E. Bernstein

#### Data and Code Availability

The raw counts and processed dataset of both the discovery and validation cohort are available at the single-cell portal with study ID: SCP2389 at: https://singlecell.broadinstitute.org/single_cell/study/SCP2389/programs-origins-and-niches-of-immunomodulatory-myeloid-cells-in-human-gliomas

Scripts and codes used to generate all the data in the study are available at: https://github.com/BernsteinLab/Myeloid-Glioma

An online tool to calculate usages of the presented consensus myeloid programs for glioma-associated myeloid cells from other experiments can be found at: https://consensus-myeloid-program-calculator.shinyapps.io/shinyapp/

This tool enables users to upload their own gene expression matrix from scRNA-seq data and output consensus program usages for each cell.

#### Human Subjects

Adult male and female patients at Massachusetts General Hospital or Brigham and Women’s Hospital (MGB) provided preoperative informed consent to take part in the study in all cases under the approved Institutional Review Board Protocol DF/HCC 10-417. Patients’ clinical characteristics are summarized in (Table S1). Patients in other cohorts were consented according to their published methods^1–3^. Previously unpublished patient data from McGill University was collected as reported with other tumors from McGill University^4^.

#### Primary tumor processing for Seq-Well and glioma organoids (GBOs)

Fresh tumor samples were collected directly from the operating room at the time of surgery and presence of glioblastoma was confirmed by frozen section. Samples were dissected into small pieces and mixed. For samples with enough material, we divided the mixed tumor pieces, with part of them going towards single cell dissociation and part going towards GBO generation.

##### Single cell dissociation and Seq-Well prep

For the MGB cohort, minced tissue pieces were mechanically and enzymatically dissociated using the Tumor Dissociation Kit, human according to manufacturer instructions and the GentleMACS™ Octo Dissociator with Heaters (Miltenyi Biotec) using custom settings. The single cell suspension was then depleted of dead cells and debris using magnetic-activated cell sorting (MACS, Dead Cell Depletion Kit, Miltenyi Biotec). Cells were then distributed drop-wise onto a Seq-Well microwell array preloaded with mRNA capture beads and processed as described previously^5^. For the other cohorts, the samples were processed as previously described^1–3^.

##### Creation and maintenance of GBOs

Minced tissue pieces were further dissected using two scalpel until tissue pieces were 1-2 mm in diameter. These were washed, further processed, and maintained according to the detailed protocol by Jacob et. al.^6^.

#### Patient PBMC/Monocyte processing

Patient PBMCs were collected at the time of surgery and isolated using SepMate-15 tubes (StemCell Technologies) and Lympholyte-H (Cedarlane) according to manufacturer’s instructions. Cells were either directly processed for Seq-Well as above, or were enriched for pan-myeloid cells using CD11b beads (Miltenyi Biotec, Cat#: 130-097-142) on Miltenyi magnet according to the manufacturing protocols and then processed for Seq-Well. CD11b+CD45+ purity was checked by flow cytometry (purity>90%).

### GBO perturbation and single-cell read-out methods

#### GBO perturbations

For perturbation experiments, GBOs were pipetted into ultra-low adherence round-bottom 96-well plates (Corning #7007) at 1 GBO per well. GBOs were plated in 100 uL of GBO media. Small molecules were then added in an additional 100 uL of media at 2x concentration. Media was changed every 2-3 days by removing 100 uL and replacing it with 100 uL of fresh media with the perturbation. Depending on the experiment, each condition had 6-12 GBOs per condition to account for heterogeneity among GBOs. For experiments with flow cytometry or scRNA-seq as a read out, multiple GBOs were grouped together in replicates per condition and then dissociated to single cells together.

#### Myeloid-GBO co-culture

Human CD11b+CD45+ cells isolated from tumor or donor patient PBMCs, as described above, were aliquoted and frozen down at 5×10^6-1×10^7 cells per ml per vial. Before co-culturing with GBOs, myeloid cells were gently thawed and washed in warm myeloid cell media (ImmunoCult™-SF Macrophage Differentiation Medium - using base media with only, M-CSF 50 ng/mL | STEMCELL Technologies, Cat. 10961), and plated in a 24-well low-attachment plate (Corning) to recover for 30 minutes in the 37C CO2 incubator. Plates were placed on an orbital rotator at 120 rpm with 2.5 x 10^6 maximum cells per well to avoid cell attachment and to maintain monocyte morphology. 10,000-50,000 monocytes, depending on the experiment, were then added to each GBO well in 100 uL myeloid cell media with a small molecule perturbation when applicable. Media was changed every 2-3 days by removing 100 uL and replacing it with 100 uL of fresh media (1:1 mix of GBO media and myeloid cell media) with the perturbation when applicable.

#### Dissociation of GBOs

In brief, all GBOs within each experimental replicate were grouped together in a 1.7 mL Eppendorf tube, media was aspirated, and GBOs were washed two times with 1 mL media to remove small molecules and/or cells. GBOs were then dissociated to single cells using dissociation media from the Miltenyi tumor dissociation kit mixed 2:1 with Accutase in the 1.7 mL tubes. These were placed at 37C and the mechanically dissociated every 5-10 min via pipetting up and down until there was a homogeneous single cell mixture. Cells were passed through a 40 um filter and then used for downstream assays. Cells were processed for SeqWell as described above or analyzed by flow cytometry as described below.

#### Flow cytometry

Flow cytometry was done based on a prior protocol^7^. In brief, an antibody cocktail was made by 1:1 ratio of Brilliant Stain Buffer(BD Horizon,566349) and PBS, and then antibodies/dyes were added. Single cell suspensions were washed by PBS+0.5% BSA in 1.5ml Eppendorf tubes or 96-well U bottom plates. Cells were pelleted by centrifugation at room temperature, 300 x g for 5 mins. After removing the washing buffer, cells were resuspended in 100ul staining cocktails via pipetting up and down ∼10 times. Plates or tubes were covered to avoid light and stained in a dark at room temperature for 20-25 mins. Cells were washed by PBS+0.5%BSA, centrifuged at room temperature, 300 x g for 5 mins. Cell pellets were resuspended in 200 ul of PBS+0.5%BSA. Flow cytometry was processed on BD LSRFortessa X-20 according to the manufacturing procedure. UltraComp eBeads(Invitrogen, 01-3333-42) are used to pre-annotate the compensation. FlowJo V10 is used to process data analysis. Antibodies used listed below.

#### Histological assessment of GBO experiments

GBOs were fixed in 4% formaldehyde for 30 min and then washed with DPBS and left in a 30% sucrose solution overnight to dehydrate the tissue. The organoids were then embedded in OCT (Sakura Tissue Tek) and frozen via isopropanol bath. The tissue was then sectioned at a thickness of 8 um using a Cryostat. For staining, slides were dried at room temperature for 10 min then prewashed with 1X TBST to remove OCT. The tissue was blocked with a glycine BSA solution for 1h at room temperature. Tissue sections were then incubated with primary antibodies (see below) either at 4C overnight or at room temperature for 2h. Tissue sections were then washed thoroughly with 1X TBST and incubated with fluorophore conjugated secondary antibodies and DAPI for 1h at room temperature. The tissue was washed with 1X TBS and incubated with 1X True Black Autofluorescence quencher for 1 min. Tissue sections were washed with 1X TBS and mounted with Prolonged Gold mounting media (Invitrogen) and covered with glass slips. For imaging, the Leica Thunder microscopy system was used with an automated mechanized stage. Images were taken using the scanning features with a 40X oil immersion objective. Images were then stitched together and enhanced with the fast computational clearing programs of the Leica LAS X software.

#### Histological Analysis

All histological images were analyzed using the Qupath open source image analysis software. The cells were counted using the cell detection feature using the DAPI channel. The detected cells were then called for positivity of up to three fluorescent markers using the single object measurement feature with positivity thresholds adjusted on a per experiment basis. Thresholds were set by comparison of experimental conditions to control and then applied to all images of the experiment through automated scripts.

| Antibodies | Source | Identifier |
| --- | --- | --- |
| Anti-SOX2 antibody [20G5] (ab171380) | Abcam | Ab171380<br>RRID:AB_2732072 |
| Anti Iba1, Rabbit (for Immunocytochemistry) | Fujifilm Wako | 019-19741<br>RRID:AB_839504 |
| CD3 epsilon antibody [CD3-12] | Genetex | GTX11089<br>RRID:AB_369097 |
| Cleaved Caspase-3 (Asp175) Antibody #9661 | Cell signaling | 9661<br>RRID:AB_2341188 |
| P2Y12 (extracellular) Polyclonal Antibody | Thermo Fisher scientific | PA5-111827<br>RRID:AB_2857236 |
| CD45 Monoclonal Antibody (YAML501.4) | Thermo Fisher scientific | MA5-17687<br>RRID:AB_2539077 |
| TMEM119 Polyclonal antibody | Proteintech | 27585-1-AP<br>RRID:AB_2880915 |
| Anti-EGFR (Ab-1) Mouse mAb (528) | Milliporesigma |  |
| Recombinant Anti-VSIG4 antibody [EPR22576-70] (ab252933) | Abcam | Ab252933 |
| CD163 mouse monoclonal antibody, clone 2G12 | Origene | 50-167-6602<br>RRID:AB_2623740 |
| Anti-Ki67 antibody | Abcam | Ab15580<br>RRID:AB_443209 |
| CD68 Monoclonal Antibody (KP1) | Thermo | 14-0688-82<br>RRID:AB_11151139 |
| MRC1(CD206) antibody | Biolegend | 321102<br>RRID:AB_571923 |
| NRP1 (CD304) Antibody | Thermo | R F : 2876<><br>WWNI FGd7=5>9 ; < |
| Nurr1 antibody | Thermo | PA5-78097<br>RRID:AB_2736269 |
| CD83 Antibody | Biolegend | 305302<br>RRID:AB_314510 |
| RHOB antibody | Thermo | 711274 |
|  |  | RRID:AB_2633147 |
| CXCR4(CD184) Antibody | Biolegend | 306502<br>RRID:AB_314608 |
| P2RY12 Antibody-BV421 | Biolegend | 392106 |
| CD8 Antibody-BV711 | Biolegend | 344374 |
| TMEM119 Antibody-AF488 | Abcam | AB225497 |
| CD163 Antibody-PE | Biolegend | 333606 |
| CD11b Antibody-PE-Cy5 | Biolegend | 301308 |
| CD45 Antibody-AF700 | Biolegend | 304012 |
| ICAM1(CD54) Antibody-PacBlue | Biolegend | 332716 |
| CD16 Antibody-BV711 | Biolegend | 302044 |
| **Flow Cytometry Reagents** |  |  |
| UltraComp eBeads | Invitrogen | 01-3333-42 |
| Brilliant Stain Buffer | BD Horizon | 566349 |
| Zombie NIR Live/Dead Dye | Biolegend | 77184 |

### Single-cell, spatial, and bulk RNA-seq analyses

#### Cohorts

There were 4 cohorts utilized in this study, split into two datasets. The discovery dataset contained the MGB, Houston Methodist^1^, and Jackson Laboratories^3^ cohorts. Cells in these cohorts were assayed with more advanced scRNA-seq technologies: Seq-Well S^3^ (MGB) or 10X Genomics 3’ v3 (Methodist and Jax Labs). The validation dataset was composed of samples from the McGill cohort, some of which had been previously published ^2^;^4^ and a set that were not previously published. McGill tumors were assayed using 10X Genomics 3’ v2 kits.

#### Alignment

The Cumulus platform^8^ was utilized to handle the processing of the large-scale Single-cell RNA-Seq experiments. Libraries were aligned to the GRCh38 genome using STARsolo^9^. See our GitHub page for specific settings https://github.com/BernsteinLab/Myeloid-Glioma.

We merged the STARsolo raw outputs (i.e., no filtration of cells) into a single expression matrix per study using Seurat’s “Read10X” and “merge” functions. We removed cells in which the expression of less than 500 genes or more than 6000 genes was detected. We also filtered out cells that demonstrated less than 1000 UMIs. We have also removed genes expressed in less than three cells in the matrix.

#### Data processing and visualization

The raw matrices outputs of STARsolo for each tumor were gzipped and used as input for Seurat ^10^ by utilizing the Read10X() function with the default parameters. The pipeline was performed for each cohort independently. Tumors belonging to each cohort were merged using Seurat’s merge() function to generate a Seurat object for each cohort. The percentage of mitochondrial gene expression was determined using PercentageFeatureSet() with the pattern set to “^MT.”. We filtered out cells expressing below 500 genes and above 6000 genes. We also filtered out cells with less than 1000 UMIs and cells with more than 25% of transcriptome composed of mitochondrial gene expression. The filtering process was carried out using Seurat’s subset() function.

For plotting purposes, normalization, scaling, and variable gene detection were performed using the SCTransform() function, where we used the percentage of mitochondrial gene expression as a regression factor. We performed PCA using RunPCA() with default parameters and generated an elbow plot using the ElbowPlot() function to help us determine the dimensions for generating UMAPs and for Louvain clustering (MGB: 24, Houston Methodist: 19, Jackson’s Laboratory: 16).24

UMAP was generated using the RunUMAP() with the reduction set to “pca”. FindNeighbors() and FindClusters() were used for clustering, with the resolution set to 0.3.

#### Classification of tumor cell types

To classify tumor cells in all cohorts, we identified the main cell programs in the MGB cohort and identified the top program for each cell in all cohorts. This top program was then used as the cell’s classification.

We merged all cells from the 22 tumors in the MGB cohort and used this expression matrix as the input for cNMF. We identified the top 4000 most variable genes using SCTransform, regressing out mitochondrial content. We subsetted the matrix for these genes and the resulting matrix was then subjected to consensus non-negative matrix factorization (cNMF) ^11^.

For the cNMF “prepare” function, we performed factorization over K ranges from 2-35. We ensured that all the variable genes were considered for the factorization using the parameter “-- numgenes 4000”. We also performed 500 iterations by inputting “--n-iter 500” in the cNMF prepare script. K=18 was the highest value with silhouette score above k=5 and was thus chosen for the “consensus” script of cNMF. cNMF was run with “--local-density-threshold” value at 0.015.

We annotated each program on the final “gene_spectra” output of cNMF by comparing the top 100 genes to previously published gene sets and known marker genes. gProfiler ^12^ was used to determine enrichment scores for a manually curated gene set matrix with over 600 gene sets (Table 3). Manual integration of enrichment scores and known marker genes helped us determine the names of the programs (Extended Data Figure 1a). Of note, MGH720, a tumor with histological diagnosis of Giant Cell Glioblastoma, had a cNMF unique malignant program.

We then used the gene spectra output of the cNMF programs to calculate the usages of these programs by cells in the other published cohorts. We extracted a raw counts matrix including the intersection between genes detected in the cohort and the top 4000 variable genes in the MGB cohort. This matrix was then subjected to the cNMF “prepare” script for normalization. The -- numgenes parameter is set to the number of genes in the matrix. We used sklearn.decomposition.non_negative_factorization in which X is the filtered normalized expression matrix, and H is the filtered gene spectra consensus matrix. The following parameters were used: “n_components= 18, init=’random’, update_H=False, solver=’cd’, beta_loss=’frobenius’, tol=0.0001, max_iter=1000, alpha=0.0, alpha_W=0.0, alpha_H=’same’, l1_ratio=0.0, regularization=None, random_state=None, verbose=0, shuffle=False”. The code is available at https://github.com/BernsteinLab/Myeloid-Glioma.

Finally, each cell was annotated as a cell type using the final “usage” matrix output of cNMF or the calculated usage matrices as discussed above. The usage scores were normalized to 100% for each cell. For each cell, the usage scores for all programs in each category were summed to create a usage score for the cell type category. For example, the usage scores for 4 myeloid programs were summed to create the “myeloid usage” per cell. Cells were then annotated as one of the cell types using the top scoring usage for cell type category.

Of note, cycling cells were considered separately. inferCNV was used to annotate cycling cells as “Malignant” or “Non-Malignant”. Non-Malignant cells were then additionally annotated by the next highest cell type. These secondary annotations were used when separating cell types for further cell-type specific analysis.

#### CNA inference from single-cell data

We selected a group of reference cells not annotated as any malignant program from various tumors (i.e., a mix of Myeloid, T cells, Oligos, and Vasculature Cells). We extracted and merged the raw counts of these reference cells into a single matrix. The reference cells used are given in (https://github.com/BernsteinLab/Myeloid-Glioma). We then utilized the inferCNV package (inferCNV of the Trinity CTAT Project. https://github.com/broadinstitute/inferCNV). We performed the analysis for each tumor separately. In the annotation file, we included the reference cells and annotated the cells of each tumor, as discussed above. We concatenated each tumor’s raw matrix with the reference cells’ raw matrix. We constructed the gene order file required for inferCNV using the “gtf_to_position_file.py” script provided by the inferCNV package. We have included the following additional arguments: “--denoise --HMM --cluster_by_groups --cutoff 0.1”. We have also ensured that the --ref_group_names match the names given to the reference cells in the annotation files. The selection of the reference cells was performed for each cohort separately.

#### Doublet Detection

Doublets were determined using integration of cNMF and inferCNV data. Cells were considered Doublets by cNMF if they expressed a second program above a specific threshold. Cell-type-specific thresholds were selected by subsetting by cell type, then plotting the usage of each potential second program. From this plot, we found the value which separated the background usage of a second program from doublets. Cells were also considered doublets if their cNMF annotation was not compatible with the inferCNV profile. Of note, the cycling programs were not considered in doublet analysis.

#### Integrated definition of malignant cells

If a tumor had detectable CNVs by inferCNV, cells from that tumor needed to meet the following criteria: Non-doublet, positive for CNV, and not annotated as a non-malignant cell type by cNMF program. For those tumors in which CNVs could not be readily detected by inferCNV, we relied on annotations based on cNMF.

#### Gene program identifications

For more granular analysis of cell programs for a specific cell type (myeloid cells, T cells, or malignant cells), we took cells in each specific category and removed doublets based on the method described in the “Doublet Detection” section above. We then input only cells determined to be singlets into another cNMF analysis for each category.

##### Myeloid cells

We used the MGB, Jax Labs, and Methodist Cohorts for identifying the cNMF programs in myeloid cells in Gliomas. The cNMF was carried out in two rounds for each cohort. The first round was used to identify cells using programs that are not myeloid (i.e., different cell type identity) or programs used by less than 100 myeloid cells. We remove such cells for subsequent analyses. The second round was used to determine the myeloid programs (Supplemental Table 2).

In the first round, raw counts of all cells annotated as myeloid and singlets (non-doublets) from each cohort were used to create a Seurat object independently. We then normalized the Seurat object using NormalizeData() and identified the top 2000 variable genes with mean expression above 0.001 in expressing cells in each cohort using the FindVariableFeatures(). Subsequently, we output the three matrices. These matrices were subjected separately to cNMF with the following parameters in the “prepare” script: --n-iter 500 --total-workers 1 -- seed 14 --numgenes 2000. Then we performed factorization and generated the K-plots using the factorize, combine, and k_selection_plot scripts of cNMF. We then chose the following Ks: MGB - 22, Houston Methodist - 23, Jackson’s laboratories - 14. We then performed the consensus script with the above Ks and a “local-density-threshold” of 0.02.

In the second round, we removed cells from each cohort as discussed above and we created a merged Seurat object from the three cleaned matrices using Seurat’s merge() function. Then, we normalized the merged Seurat object and detected variable genes using NormalizeData() and FindVariableFeatures(). We then filtered out the genes with a mean expression value below 0.01 in expressing cells and standardized variance below 1. We then filtered the cleaned myeloid matrix of each cohort to include the variable genes that met the criteria mentioned above. Similar to round 1, these matrices were subjected to cNMF individually with the following parameters in the prepare script: --n-iter 500 --total-workers 1 - -seed 14 --numgenes 2276. Then, we also ran the factorization and generated the K-plots using the factorize, combine, and k_selection_plot scripts of cNMF.We then chose the following Ks in the second round: MGB - 18 (We filtered out programs that are not myeloid), Houston Methodist - 19, Jackson’s laboratories - 18. Finally, we then performed the consensus script with the above-mentioned Ks and a “local-density-threshold” of 0.02.

To find the consensus programs, we performed a cosine correlation of the gene spectra output of each cohort. Programs with a cosine similarity score of 0.5 or above were considered for further processing. These programs’ weights ‘w’ were then averaged to obtain a set of meta-programs representing the shared transcriptional programs across datasets. Ward’s method, a hierarchical clustering algorithm, was applied to the similarity matrix to visualize the relationships between programs in a heatmap.

We averaged the spectra scores in the “gene_spectra_consensus” outputs of round 2 cNMF for programs with high cosine similarity, resulting in 14 consensus myeloid programs across the three cohorts. We annotated the programs as discussed above.

##### Malignant and T cells

Malignant cells and T cell programs (Supplemental Table 2) were obtained from the MGB data in separate cNMF runs similar to the two-step cNMF used in myeloid cells. We selected a k-value of seven for the malignant cells based on the silhouette plot’s stability, consistent with previously published glioblastoma signatures represented in our five chosen programs ^10^. For the T cells, we found the optimal program count to be four. We calculated the usage of these programs in the other cohorts in a way similar to the all-cell type cNMF mentioned above.

#### Processing and cNMF for PBMC scRNA-Seq libraries

The PBMC libraries were processed for cNMF similarly to the primary tumor libraries. We merged the expression matrix of all the PBMC libraries using Seurat’s “merge” function. The seurat object was then normalized using’ NormalizeData()” and “ScaleData()”. We then used “FindVariableFeatures() “to calculate the variance score for every gene. We selected the top 3000 variable genes after removing genes below 0.001 mean expression (in expressing cells) and then subsetted the gene expression matrix to include the variable genes only. As described above, cNMF was performed with “--numgenes 3000” and the value K=18 for the “consensus” script of cNMF, annotation was done using gProfiler, and non-doublet cells were identified. We isolated myeloid cells, identified the top 2000 most variable genes, and performed two rounds of cNMF (K=16)

#### Comparison of gene programs

To assess the similarity of two given gene programs, we took the top 100 genes in those programs and compared their makeup using Jaccard index. P-values were measured by assessing the probability of observed gene matches were obtained by random chance using a binomial test where k is the number of matches, n is the size of the gene set, and p is probability of randomly drawing matches from all genes scored in the program.

#### Comparative UMAP and Clustering of myeloid cells

We extracted the raw counts of all MGB cells annotated as myeloid and singlets (non-doublets) from each tumor. Then, normalization was performed for each tumor separately using NormalizeData() with default settings, followed by FindVariableGenes() with the following settings (selection.method= “vst”, nfeatures = 2000). We then ran FindIntegrationAnchors() k.filter set at 30 (to ensure that tumors with few myeloid cells were included. We then used the anchors identified as input to batch-correct the objects using IntegrateData(), setting features.to.integrate as the intersection of genes detected in all tumors in and dims to 1:30.

The batch-corrected Seurat object was then subjected to ScaleData(), RunPCA(), and ElbowPlot() with default parameters to identify the number of dimensions to use for Louvain clustering and UMAP generation. We generated the UMAP using RunUMAP() with “dims” set to 1:8 and “reductions” set to “pca”. We performed the clustering using FindNeighbors() with “dims” set to “1:8” followed by FindClusters() with a 0.3 resolution. UMAPs were generated using the “DimPlot()” function.

#### Generation of heatmap for gene expression programs

To generate the gene expression heatmap of the NMF programs, we assigned the myeloid cells to one of the following categories:

*Microglia*: Minimum 10% usage of microglia program and other identity programs are all below the usage value of the microglia program (macrophages must be below 10%).

*Microglia-Like* - Minimum 10% usage of microglia and 10% usage of monocytes or macrophages program. Other identity programs should be below the usage value of these two programs (Otherwise, it is assigned as a microglia).

*Macrophages* - Minimum 10% usage of macrophage program and other identity programs are all below the usage value of the macrophage program (monocyte below 10%).

*Mono_Macro* - Minimum 10% usage of macrophages and 10% usage of monocytes program. Other identity programs are below the usage value of these two programs.

*Monocytes*: Minimum 10% usage of macrophage program and other identity programs are below the monocytes program’s usage value.

*cDC* - Minimum 10% usage of the cDCs program and other identity programs are all below the usage value of the cDCs program.

*Neutrophils* **-** Minimum 10% usage of the Neutrophils program and other identity programs are all below the usage value of the Neutrophils program.

*Activity Dominated* - All identity programs are below 10% usage.

For selecting the genes included in the heatmaps, we identified the top 100 genes in the averaged gene_spectra output of the myeloid cNMF programs for each program. We counted the number of myeloid cells expressing the top 100 genes in each program. We included the top 20 genes with the highest number of myeloid cells expressing them in each program.

We used the “ComplexHeatmap” library in R ^13^ to generate the heatmaps. We z-score scaled the log1p normalized gene expression values across all the myeloid cells (regardless of the categorization). We then set an upper limit of 2 and a lower limit of −1. We generated a heatmap for each category separately. We turned off row clustering (genes) by setting “cluster_rows = FALSE” in the Heatmap function, and we allowed default column clustering within each category (cells).

#### Generation of quadrant plots

The X-axis of the quadrant plots is calculated by subtracting the usage of the C1Q Immunosuppressive program from the IL1B pro-inflammatory program. In contrast, the Y axis is calculated by subtracting the Scavenger Immunosuppressive program usage from the CXCR4 pro-inflammatory program in each myeloid cell. For the quadrant plot with scatterpies as dots, we used the “scatterpie” library (https://github.com/GuangchuangYu/scatterpie). The “others” category for the pie charts was calculated by summing the usages of all programs excluding the four immunomodulatory programs.

#### Assignment of myeloid and T cells to recurrent gene program

We considered the myeloid cell to use an activity cNMF program if it has a minimum usage of 20%. For identities, we annotated the myeloid cells with the top identity program usage if it has at least 10% in a particular identity program. A single myeloid cell could be classified as using multiple programs. For example, a myeloid cell can be considered microglia using the Scavenger Immunosuppressive program if it has at least 10% usage of microglia program and 20% usage of the scavenger immunosuppressive program. For Extended Figure 4b-c, we calculated Observed/Expected ratios of the co-occurrence of a myeloid identity program and a myeloid activity program and used a hypergeometric test to assess significance.

T cell program usages were more distinct. We therefore simply defined them by the program of their top usage.

#### Creation of discretized matrix and identification of marker genes per program

To facilitate the discovery of specific markers and the spatial cellular demultiplexing described below, we created a discretized matrix of MGB cells with the strongest expression of each tumor cell program, including each myeloid, T cell, and cancer programs, thus excluding intermediate cells. For the outputs of Myeloid, Malignant, and T cell cNMFs, cells with a minimum 2.5-fold higher usage of a particular program over the second most used program were annotated with that program as a discrete cell. For oligodendrocytes and vasculature, the usages from “all cell types” cNMF outputs were used to annotate Oligo or vasculature discrete cells. Doublets, cycling programs, and cycling cells were excluded from the analysis.

Additionally, we downloaded scRNA-Seq for normal brain tissue from 25 and 40-year-old individuals ^14^. These libraries were processed using the abovementioned Seurat processing; we normalized the cells and used 1:14 as “dims” for generating UMAPs and identifying clusters. We performed FindMarkers and extracted neurons and astrocytes from the published matrix based on differentially expressed genes. We then merged these cells with the discrete cells matrix. We generated a UMAP as described above.

To identify markers for the immunomodulatory programs, we extracted discrete cells annotated as Scavenger Immunosuppressive, C1Q Immunosuppressive, IL1B inflammatory, or CXCR4 inflammatory using Seurat’s subset() function. A similar processing pipeline was performed with the option “dims” set to 1:16 in FindNeighbors() and FindClusters(). The UMAP coordinates for these cells were obtained using Embeddings() with the option “reduction” set to “umap”. Then the normalized matrix of these cells was extracted using the function GetAssayData() with the option slot set to “data”. These files were used as input for COMET ^15^ to identify the significant markers that distinguish each immunomodulatory program.

#### MAESTER analysis for determination of myeloid cell origin

To determine the origins of the various myeloid cell identities, we processed the MAESTER libraries in three steps, namely, (1) preprocessing, (2) identifying variants of interest, and (3) measuring the enrichment of the identified variants in the various myeloid identities.

*Step 1: Preprocessing.*The preprocessing was performed as previously described^16^. Briefly, we trimmed high quality reads, aligned them using STAR (hg38), and processed them using MAEGATK as previously described^16^.

*Step 2: Low-resolution pseudobulking to identify variants of interest.* To determine the origin of the myeloid cell identities, we had to identify variants specific to myeloid cells in the tumor microenvironment and not present in the PBMC. We also had to identify variants present in the myeloid cells in the PBMC but absent in the tumor microenvironment. We pseudobulked the primary tumor libraries to the following categories based on their RNA-Seq annotation:

For primary tumor libraries: “Malignant”, “Tumor-Associated Myeloid (TAMs)”, “Stromal”, “Oligo”, “Tcells” and for PBMC libraries, we considered only “Neutrophils” and “Monocytes” (Myeloid_PBMCs).

This stage involves multiple steps:

1- The first step was to extract the number of UMIs supporting each possible variant at every possible location from MAEGATK output using the following script in R:

~~~
computeAFMutMatrix <- function(SE){
 ref_allele <- as.character(rowRanges(SE)$refAllele)
getMutMatrix <- function(letter){
 mat <- (assays(SE)[[paste0(letter, “_counts_fw”)]] + assays(SE)[[paste0(letter, “_counts_rev”)]])
 rownames(mat) <- paste0(as.character(1:dim(mat)[1]), “_”, toupper(ref_allele), “>”, letter)
 return(mat[toupper(ref_allele) != letter,])
}
rbind(getMutMatrix(“A”), getMutMatrix(“C”), getMutMatrix(“G”), getMutMatrix(“T”))
}
~~~

The computeAFMutMatrix generates a matrix where the rows are every possible variant in the mitochondrial genome, and the columns are barcodes (cells). The values represent the UMIs supporting each variant in the given barcode.

2- We extracted the coverage of the Maester libraries at each base of the MT genome in every cell from the output of MAEGATK, which is stored in assays(maegatk.rse)[[“coverage”]] whereby the “maegatk.rse” is the R object output of MAEGATK.

3- We annotated the cells in each matrix (obtained from step 1 and step 2) based on the scRNA-Seq library (as mentioned in the ***“Annotation of cells to broad cell type categories”***). We created a data frame in which one column has the barcodes, and the other has the respective annotation.

4- We pseudobulked the UMI count matrix (obtained from Step 1) using R using the following steps: (a) we subsetted the matrix into each annotation using tibble. (b) Then, we used the sum() function to sum all the rows in each matrix, creating a pseudobulked number for each annotation. (c) We merged all the summed values in each matrix for each variant possibility into a pseudobulked matrix in which each column represents an annotation.

5- We pseudobulked the coverage matrix (obtained from Step 2) similarly to Step 4.

6- We calculated the pseudobulked Variant Allele Frequencies (VAFs) using R. We added a pseudo-count 0.000001 to each value in the pseudobulked coverage matrix. We divided each value in the pseudobulked counts matrix by its respective coverage in the pseudobulked coverage matrix to obtain pseudobulked VAFs for each category.

7- To ensure the specificity of each variant’s detection with 99% certainty, we utilized a binomial model to establish a minimum VAF threshold dependent on coverage.

8- To consider the variant to be specific to the myeloid cells in the tumor microenvironment, the variant has to meet the following criteria: (a) meet the minimum VAF requirement for TAMs for the coverages in TAMs and Myeloid_PBMC categories. (b) VAF=0 in the Myeloid PBMC category. (c) VAF > minimum required in the TAM category.

9- To identify variants specific to the myeloid cells in PBMC, the variant has to meet the following criteria: (a) meet the minimum VAF requirement for Myeloid_PBMC for the coverages in Malignant, TAMs, and Myeloid_PBMC categories. (b) VAF = 0 in the Malignant category. (If the tumor library is enriched for malignant cells, this criteria can be replaced with VAF in Myeloid_PBMC, which is 20 times more than Malignant (c) VAF > 0 in the TAM category. (d) VAF > minimum required in Myeloid_PBMC category.

*Step 3: Determination of enrichment of variants of interest in various identities of myeloid cells.* We hypothesized that myeloid cells enriched for tumor microenvironment-specific variants are tissue residents, whereas those enriched for PBMC-specific variants are monocytes-derived. To perform the enrichment analysis, the following steps were implemented:

1- We annotated the TAMs with the myeloid identities cNMF programs. We kept cells that met the following criteria to ensure the reliability of the identity and the results:

For Microglia annotation - (a) Cell has a minimum of 15% usage of microglia program. (b) the microglia program must be at least two times higher than any other identity program. (c) macrophage program has to be at least two times lower than any other identity program.

For Microglia_Like annotation - (a) Cell has a minimum of 10% usage of microglia program. (b) The cell has a minimum of 10% usage of the macrophage program. (c) the microglia program must be at least twice as high as any other identity program (excluding macrophages and monocytes). (d) The macrophage program must be at least twice as high as any other identity program (excluding microglia and monocytes).

For Mono_Macro annotation - (a) Cell has a minimum of 10% usage of monocytes program. (b) The cell has a minimum of 10% usage of the macrophage program. (c) the monocyte program must be at least two times higher than any other identity program (excluding macrophages). (d) The macrophage program must be at least twice as high than any other identity program (excluding monocytes).

For Macrophages annotation - (a) Cell has a minimum of 15% usage of the macrophages program. (b) the macrophage program has to be at least two times higher than any other identity program. (c) monocytes program has to be at least two times lower than any other identity program. (d) microglia program has to be at least two times lower than any other identity program.

For Monocytes annotation - (a) Cell has a minimum of 15% usage of the monocytes program. (b) the monocyte program has to be at least two times higher than any other identity program. (c) macrophage program has to be at least two times lower than any other identity program.

For Neutrophils annotation - (a) Cell has a minimum of 15% usage of neutrophils program. (b) the neutrophils program has to be at least two times higher than any other identity program.

For cDC annotation - (a) Cell has a minimum of 15% usage of cDC program. (b) the cDC program has to be at least two times higher than any other identity program.

2- We subsetted the single cells MAESTER UMI matrix obtained in (Stage 2 - Step 1) to smaller matrices, with each matrix composed of cells annotated with one of the above identities (Stage 3 - Step 1).

3- We created a matrix in which the rows are the variants of interest identified in Stage 3, and each column represents an identity. The values in this matrix denote the number of cells in each identity in which the variant was detected. We used “annotation_matrix[,apply(annotation_matrix,2,function(x) sum(x > 0))]” in R for each subsetted matrix. This script will keep columns that have UMI > 0. Then, we identified the number of cells having UMIs >0 for each annotation by using “ncol(as.matrix(annotation_matrix)” whereby annotation_matrix represents each subsetted matrix.

4- We removed any variant with 0 value in every pseudobulked category.

5- We removed any identity (column) that had less than ten total values from consideration to ensure proper enrichment results.

6- We used GSVA ^19^ by categorizing the rows of the matrix as PBMC-specific or Tumor microenvironment-specific. Since the values are integers (number of cells having UMIs supporting the variants), we set the option “kcdf” to “Poisson” in the “gsva” script.

7- We subtracted the GSVA enrichment for tumor microenvironment-specific variants from GSVA enrichment for PBMC-specific variants for each identity. Positive values indicate PBMC-origins, whereas negative values indicate tissue resident origin.

8- We calculated the prevalence of each identity in each tumor. We plotted the dot plots with the dot’s position representing the subtracted GSVA enrichment values and the size of the dot representing the prevalence using ggplot2.

9- We calculated the average % usages of the four immunomodulatory programs in each identity, labeled the remaining percentage as others in each tumor, and plotted them as stacked bar plots using ggplot2.

#### Sample level association analysis

To measure inter-state association, we determined the average usage of myeloid, T, and cancer programs in their respective cells in each sample. We utilized Spearman correlation to understand how a rise in one myeloid state corresponded to changes in cancer or T cell states, enabling us to examine how variations in one state could influence the behavior of others within the same sample.

In our state-clinical/molecular associations analysis, we applied the sample state averages derived above. These were compared across various clinical and molecular categories, such as tumor grade, IDH mutation status, and steroid usage, using the Wilcoxon rank-sum test.

To account for the confounding effect of sample hypoxia on the influence of dexamethasone on the complement immunosuppressive myeloid program, we limited this analysis to samples where the average MES2 program usage in cancer cells was below 20%.

We employed a linear least-square regression to investigate changes in average myeloid program usage across samples relative to the daily dose of dexamethasone.

To assess the myeloid and cancer program associations with T-regs, we classified each sample in the top 50% and bottom 50% expressors of T-reg and cancer/myeloid program usages and used Fisher’s Exact Test to measure this association. P-values were adjusted using False Discovery Rate (FDR).

#### Cell cycle analysis

We evaluated the relationship between the cell cycle and different myeloid cell states in each sample independently. Cells were defined as cycling if the combined usage of G1S and G2M programs exceeded 20%, and as belonging to a specific state if its usage surpassed 20%. We used Fisher’s exact test to measure this association and obtained an odds ratio. For each program, we assessed the probability that its sample distribution was higher than 1 using a one-sample t-test on the log transformed values, so they become approximately normally distributed^65^.

#### Deconvolution of TCGA, GLASS, and G-SAM and correlation with cell programs and clinical data

The analysis pipeline consisted of three steps to deconvolve published glioma bulk expression datasets. Step 2 and 3 of the pipeline were performed for each dataset separately.

*Step 1: Creating gene set for each program.*The top 50 genes were obtained for each myeloid program by ranking them in the merged gene spectra output. For T cells and Malignant Cells, we ranked the gene spectra from their respective cNMF outputs to obtain the top 50 genes for each program. For the other cell types, we used the gene spectra output of the cNMF of all cells in the tumor microenvironment. We ranked and obtained the top 50 genes for the Pericytes, Endothelial, and Oligo programs. Then, we selected genes which only appeared in the list of a single program and not in the top 100 genes of any other program.

*Step 2: Calculating Module scores using Seurat.* Raw gene counts for Glioma datasets were obtained. The matrices were then CPM-normalized using EdgeR’s DGEList(), calcNormFactors(), and cpm() functions ^20^. The CPM-normalized matrix was then log-transformed using the log1p() function of R. Afterward, Seurat objects were created using the CreateSeuratObject() function. The Seurat objects were scaled using ScaleData(). Finally, the module scores were calculated using Seurat’s AddModuleScore() function. The above-mentioned gene sets were used as input in the “features” option of AddModuleScore().

*Step 3: Normalizing the Module Score.* In order to correct for the purity differences in the published bulk glioma mRNA-expression datasets, we imputed the percentages of Malignant, Oligo, Vasculature, Myeloid, T cells, and other immune cells in the bulk expression matrices using CIBERSORTx ^21^. As input, we discretized the MGB “all cell type” matrix in a similar fashion but by utilizing the usage outputs of the “all cell type” cNMF only. The matrix was used as the “single_cell_reference” in the Fractions module of CIBERSORTx. The published bulk gene expression matrices used as “mixture” input were also CPM normalized without log-transformation. The module scores obtained above were then divided by the imputed value of their respective cell type in the CIBERSORTx results outputs.

These normalized module scores were then used to correlate with sample clinical data, including IDH mutation status and Molecular Grade. For correlation with clinical data, the normalized module scores were log10 transformed. We removed libraries with a CIBERSORTx value below 0.1 for myeloid. To correlate the module scores with the expression levels of the identified ligands in the LGG and GBM TCGA cohorts, we performed a Pearson correlation between the log-transformed normalized module score and the log-transformed normalized expression. We used the Wilcoxon Rank-Sum test to assess statistical significance, adjusting p-values using FDR.

#### Spatial transcriptomic analysis

We used Visium spatial transcriptomics to analyze 23 samples^44^. Our methodology encompassed two distinct approaches: unbiased niche discovery using cNMF; and cell state demultiplexing using RCTD^68^ and the single-cell RNA-seq programs.

In the unbiased niche discovery, we used all samples, both normal and cancer, for cNMF analysis. We selected the 1500 top spatially variable genes based on Moran’s I and ran the cNMF algorithm from k=2 to 25. The optimal k value (k=7) was determined as the highest one with a Silhouette score above 0.9. We defined the identity of niche programs through the analysis of gene expression patterns and excluded the program whose top genes were mitochondrial and ribosomal. To ensure uniformity, the usage scores were normalized to 1 in each pixel.

For cell state demultiplexing, we used the discretized matrix described above. Using this single-cell reference, RCTD was run individually on each spatial sample. Again, the scores were normalized to sum to 1 in each pixel.

To assess the link between niche and cell state, we used Spearman correlation to evaluate the relationship between normalized RCTD scores and cNMF usage for individual patient samples. In Figure 4, a cell type was classified as part of a niche if correlation held statistical significance (FDR<0.01) in over half of the samples.

In our analysis of cell state and niche-niche spatial relationships, we designed a spatial regression model to quantify cell state proximity, analyzing one sample at a time. For any pair of cell or niche states across all pixels in a sample, we regressed the presence of one state in a central pixel against the average expression of the other state in surrounding pixels. In this model, we binarized the central pixel’s state presence using a state-specific threshold determined by a knee plot (KneeLocator^69^) of cell state expression across pixels in all samples. The resulting regression coefficient indicates the association strength between the states, with the p-value demonstrating the statistical significance of this association. The constant represents the background signal of the non-central state. Extended Data Figure 7e’s network graph was plotted using the python package NetworkX (v2.0). Each edge’s size represents the ratio of the regression coefficient to the constant, providing a measure of the state association strength relative to the background signal. Edges are statistically significant (FDR < 0.01) in over 40% of the samples with an association strength above 0.1.

#### Correlation of Immunomodulatory Programs with Responders and Non-responders to Immunotherapy

We downloaded the published Seurat object of scRNA-Seq libraries of glioma patients responding or not responding to immunotherapy ^22^. Based on the published annotations, we extracted the myeloid cells from the Seurat object. We calculated the usage of the myeloid cNMF programs in the published Seurat object, as shown above. We averaged the usage of each program in each patient and measured the difference between the usages in responding vs non-responding patients, using Wilcoxon Rank-Sum tests to assess significance.

#### Survival Analysis

We merged the survival information (Survival time) and Events from the cohorts of G-SAM and GLASS. We included IDH WT GBM tumors only. We used the normalized module score values (without log transformation) for the cNMF programs. We removed duplicate patient entries, corresponding to recurrences, by keeping the values from the primary tumor only. We removed any library with a CIBERSORTx value of 0 for the myeloid lineage. We then took the samples in the top 33% of Scavenger immunosuppressive module scores and those in the top 33% of C1Q immunosuppressive module score and considered this group of samples as the “high” immunosuppressive. For the “low” immunosuppressive group, we selected samples both in the bottom 33% of libraries in terms of Scavenger immunosuppressive module scores and the bottom 33% of C1Q immunosuppressive module scores.

The library “ggsurvfit” (https://github.com/pharmaverse/ggsurvfit) was then used to generate the Kaplan-Meier survival curve. A Cox Proportional Hazard Model (https://github.com/therneau/survival) was used to determine differences in probabilities of survival.

